# Transcriptome-inferred CIN-like fields mark antigen-presentation-low, immune-cold spatial neighborhoods

**DOI:** 10.64898/2026.05.15.725574

**Authors:** Subhajit Dutta

## Abstract

Cancer immune escape is usually interpreted as either a tumor-cell-intrinsic antigen-presentation defect or an immune-cold microenvironment. How these states are spatially arranged relative to chromosomal-instability-like genotype states remains unresolved. Here, spatial transcriptomic, single-cell and clinical transcriptomic analyses show that transcriptome-inferred CIN-like fields mark antigen-presentation-low, immune-cold neighborhoods. Across 14 melanoma tissue sections, the interaction between inferred copy-number-like burden and chromosome-expression deviation was associated with reduced MHC-I antigen-presentation activity after adjustment for tumor state, immune inflammation, technical covariates and spatial coordinates. CIN-high/AP-low spots formed coherent neighborhoods enriched for AP-low neighbors but depleted of immune-high and IFN-high neighbors under block-preserving spatial nulls. Two breast cancer Visium sections reproduced AP-low neighborhood topology, while malignant melanoma single-cell RNA-seq supported a tumor-intrinsic CNA-by-chromosome-deviation association with AP repression. Patient-level TCGA analyses linked the adverse genotype-ecotype score to poor melanoma survival. These results define transcriptome-inferred CIN-like fields as spatial correlates of immune-cold antigen-presentation failure.

## Introduction

Tumors are not uniform immunological objects. A single tissue section can contain immune-infiltrated regions, immune-desert regions, stromal barriers, necrotic or hypoxic zones, and malignant compartments with different evolutionary histories. This spatial heterogeneity matters because immune recognition is a local process: antigen presentation, effector-cell access, interferon signaling and suppressive stromal programs all occur in physical neighborhoods rather than in a well-mixed bulk average 1,2,3,4. Yet many genomic models of immune escape are still formulated at the whole-sample level. They ask whether a tumor is aneuploid, has high tumor mutation burden, has HLA loss or has low cytolytic activity, but they usually do not ask where these features are positioned inside the tissue.

Chromosomal instability provides a biologically plausible bridge between tumor genotype and immune geography. Aneuploidy and arm-level copy-number alterations are pervasive in cancer, influence gene dosage, perturb cell-state programs and correlate with adverse clinical behavior 5-9. Pan-cancer analyses have shown that highly aneuploid tumors often have lower immune infiltration and reduced markers of cytotoxic activity 6,7,13. This observation suggests that large-scale genome imbalance may contribute to immune escape, but it leaves open a key spatial question: does chromosomal instability simply identify globally immune-poor tumors, or does it organize local fields in which tumor cells with high genomic imbalance occupy antigen-presentation-low, immune-cold niches?

A spatially resolved view is particularly important for antigen presentation because AP is not only a property of malignant cells but also a readout of local inflammatory context. IFN signaling can induce HLA class I and antigen-processing genes, while immune-cold compartments may show low AP simply because inflammatory pressure is absent. Conversely, tumor cells can lose AP despite immune exposure through genetic, epigenetic or signaling mechanisms. A tissue-level analysis therefore has to distinguish low AP caused by lack of immune contact from low AP caused by failure to respond to immune contact. These alternatives cannot be separated from bulk RNA-seq alone because bulk measurements average over immune-rich and immune-poor regions.

Antigen presentation is a central axis of this problem. The MHC-I pathway requires peptide generation, transport, loading and cell-surface display through a machinery that includes HLA class I molecules, B2M, TAP1, TAP2, TAPBP, immunoproteasome components and transcriptional regulators such as NLRC5 10-12,15-17. Genetic lesions in this pathway, HLA loss of heterozygosity and defects in antigen processing are established routes of immune escape 10,11,16,17. However, loss of antigen presentation in tissue can also arise indirectly through changes in malignant state, inflammatory exposure, spot composition or local cytokine environment. Distinguishing these mechanisms requires analyses that jointly measure tumor genotype, AP activity, immune programs and spatial context.

Two biological models are especially important to separate. In an IFN-exposed AP-refractory model, tumor regions are adjacent to immune and interferon-rich neighborhoods but fail to induce antigen-presentation genes. This would imply an inflammatory interface where malignant cells resist cytokine-driven AP upregulation. In an immune-cold AP-low model, CNA/CHRDEV-high regions are not IFN-rich borders but spatial interiors depleted of immune and IFN signals, with low AP reflecting a broader cold ecotype. These models have different implications. The first suggests defective sensing or transcriptional response to inflammatory pressure; the second suggests a spatial architecture in which chromosomal instability fields occupy or produce immune-excluded territories.

The challenge is compounded by the relationship between chromosomal instability and tissue composition. High inferred CNA signal often marks tumor-rich regions, and tumor-rich regions can be immune-poor for reasons unrelated to AP regulation. A convincing analysis must therefore ask whether genotype-high/AP-low fields remain distinct after accounting for tumor state, immune abundance, IFN signaling, spatial position and technical features. It must also test whether CIN-high/AP-low regions differ from AP-low regions that are not genotype-high. This is the purpose of the matched AP-low-only controls and block-preserving spatial nulls used below.

Spatial transcriptomics is suited to this distinction because it retains tissue coordinates while measuring transcriptome-wide programs 29-36. At Visium resolution, each spot can contain multiple cells, so the analysis cannot be interpreted as a single-cell lineage map without additional support. Nevertheless, spatial transcriptomics can define tissue-level neighborhoods and test whether genomic proxies, AP scores, interferon signals and immune programs are arranged non-randomly. The key is to use spatial null models, matched controls and independent single-cell validation rather than treating spot-level associations as sufficient.

Here, I analyzed public melanoma spatial transcriptomic datasets together with breast cancer Visium sections, malignant-cell single-cell RNA-seq and TCGA patient cohorts to test whether transcriptome-inferred CIN-like genotype fields mark AP-low immune-cold neighborhoods. The genotype-state axis combined an inferred CNA-like expression shift and a chromosome-expression-deviation score, creating a two-axis representation of genome-imbalance-like transcriptional structure. Throughout the manuscript, CNA, CHRDEV and CIN-high refer to expression-derived proxies; they should not be read as direct DNA copy-number calls, subclonal copy-number maps or live measurements of chromosomal mis-segregation. The primary tissue state was defined as CIN-high for CNA and chromosome deviation and AP-low for MHC-I antigen-presentation activity. The analysis then asked four linked questions: whether the CNA-by-chromosome-deviation interaction predicts lower AP, whether CIN-high/AP-low spots form spatially coherent neighborhoods, whether those neighborhoods are immune-cold or IFN-exposed, and whether the signal is supported by non-melanoma spatial data, malignant-cell single-cell data and patient-level outcome analyses.

## Results

### A spatial genotype-ecotype framework identifies CIN-high/AP-low tissue fields

The discovery analysis used 29,341 melanoma spatial transcriptomic spots from 14 tissue sections across three public GEO datasets (GSE250636, GSE271422, GSE316760) and integrated them with 8,696 breast cancer Visium spots from two independent 10x Genomics sections (see Methods for full section-level accessions). For each spot, an antigen-presentation axis summarized MHC-I processing and presentation activity, a CNA-like axis summarized inferred copy-number-like expression imbalance, and a chromosome-deviation axis summarized chromosome-level expression deviation. Immune, cytotoxic, IFN, myeloid, CAF, hypoxia, checkpoint, tumor-state, count-depth and detected-feature covariates were retained where available. This design allowed the same spatial object to be used for genotype-state, immune-state and topology analyses without reducing the tissue to a single bulk score. (Fig. 1a; Supplementary Fig. 1).

**Figure 1.**
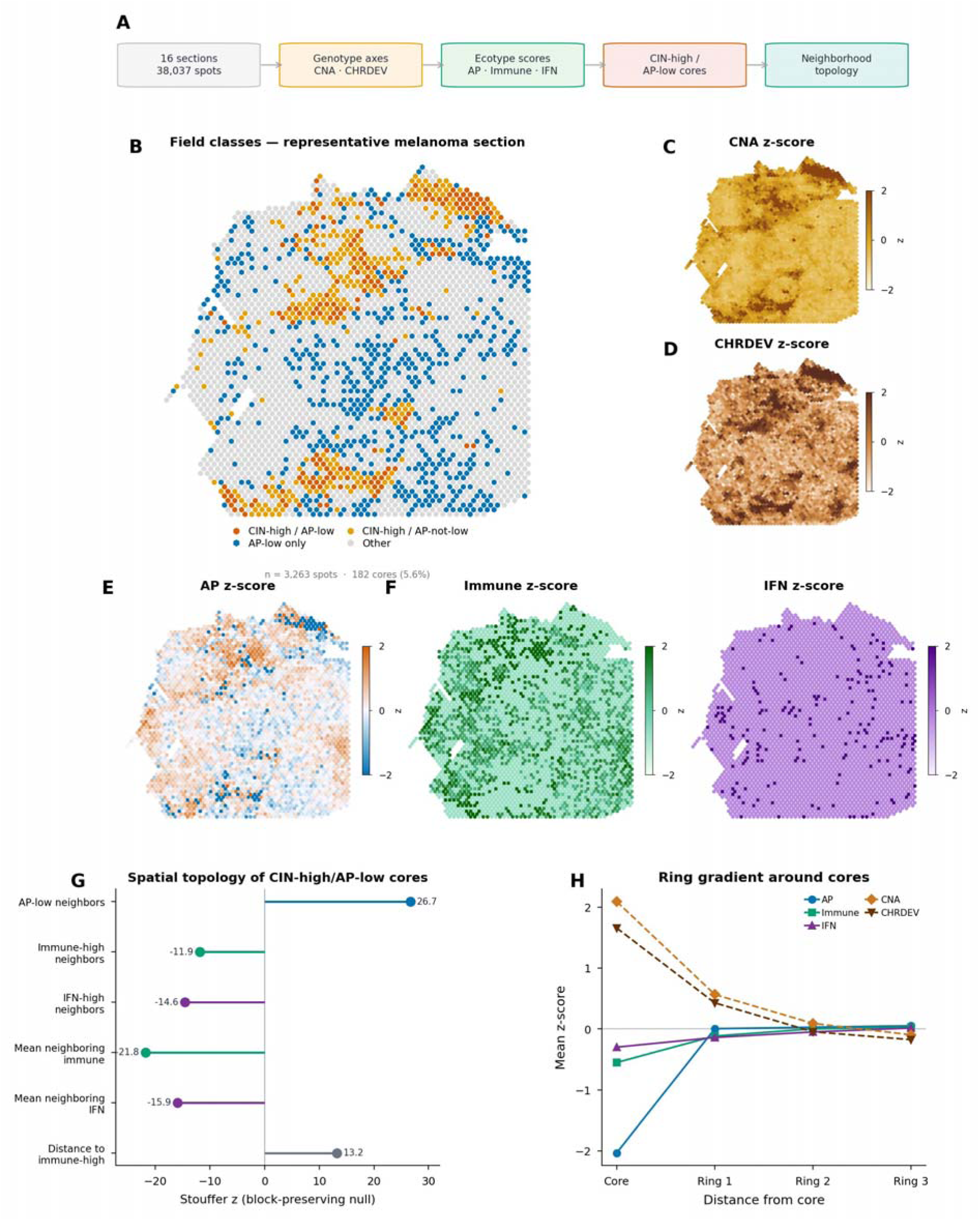
Spatial discovery of CIN-high/AP-low immune-cold fields. a, Analytical workflow from spatial sections and spots to genotype axes, ecotype scores, CIN-high/AP-low core definition and neighborhood topology. b, Representative melanoma section (GSM7983366; n = 3,263 spots; 182 cores; 5.6% prevalence) showing field classes. c,d, CNA z-score and CHRDEV z-score spatial maps. e, AP z-score spatial map with CIN-high/AP-low core outlines. f, Immune z-score and IFN z-score spatial maps. g, Block-preserving null topology across 16 sections, showing AP-low neighbor enrichment, immune/IFN neighbor depletion and greater distance from immune-high regions. h, Ring-gradient profile around CIN-high/AP-low cores for AP, immune, IFN, CNA and CHRDEV programs.

CIN-high/AP-low cores were defined within each tissue section as spots above the section-specific upper quartile for both CNA and chromosome deviation and below the lower quartile for AP. This relative definition was chosen to avoid imposing a global threshold across datasets with different chemistries, coverage and tissue processing. It yielded analyzable cores in all 14 melanoma sections and both breast sections, with section-level core counts ranging from small but sufficient focal fields to larger spatial domains. The analysis therefore focused on within-section contrasts and meta-analysis across sections rather than direct comparison of unstandardized expression values. (Fig. 1b; Supplementary Fig. 2).

The field definition was intentionally conservative. AP-low alone could arise from tissue quality, low RNA content, poor tumor cellularity or immune composition. CNA-high alone could identify proliferative or tumor-rich regions without necessarily defining antigen-presentation failure. The CIN-high/AP-low class required simultaneous genotype imbalance and AP attenuation, and all downstream topology tests compared these cores against spatially matched or block-preserving nulls.

Across the final classified spatial table, all 16 sections had at least five CIN-high/AP-low core spots and sufficient background spots for topology testing. The melanoma sections varied substantially in size and immune context, with core counts spanning focal patches and larger domains. The two breast sections had larger CIN-high/AP-low core counts, providing a useful stress test for whether the topology results were an artifact of melanoma-specific sampling. This heterogeneity was preserved in the analysis rather than normalized away; each section contributed its own effect estimate to the meta-analysis. (Supplementary Fig. 1).

The framework also separates three related but distinct quantities. The AP axis captures a tumor-recognition program, the immune and IFN axes capture local inflammatory context, and the CNA/chromosome-deviation axes capture transcriptome-inferred genotype-state imbalance. Treating these quantities separately avoids a common interpretive shortcut in which low immune signal, low AP and high tumor content are collapsed into a single immune-desert label. The analysis instead asks whether CNA/CHRDEV-high regions have a reproducible AP-low and immune-cold architecture after these axes are modeled independently.

### CNA and chromosome-deviation jointly predict AP attenuation in melanoma spatial sections

The primary melanoma model tested AP activity as a function of the transcriptome-inferred CNA-like axis, chromosome deviation and their interaction within each spatial section, with adjustment for tumor state, immune inflammation, cytotoxicity, IFN signaling, stromal/myeloid programs, sequencing-depth covariates and spatial coordinate terms. The interaction between CNA and chromosome deviation was consistently negative for AP across all 14 melanoma sections. Random-effects meta-analysis gave beta = −0.0775, 95% CI −0.1089 to −0.0462, p = 1.27 x 10^-6 and FDR = 1.71 x 10^-5, with 14 of 14 sections in the negative direction. (Fig. 2a-c).

**Figure 2.**
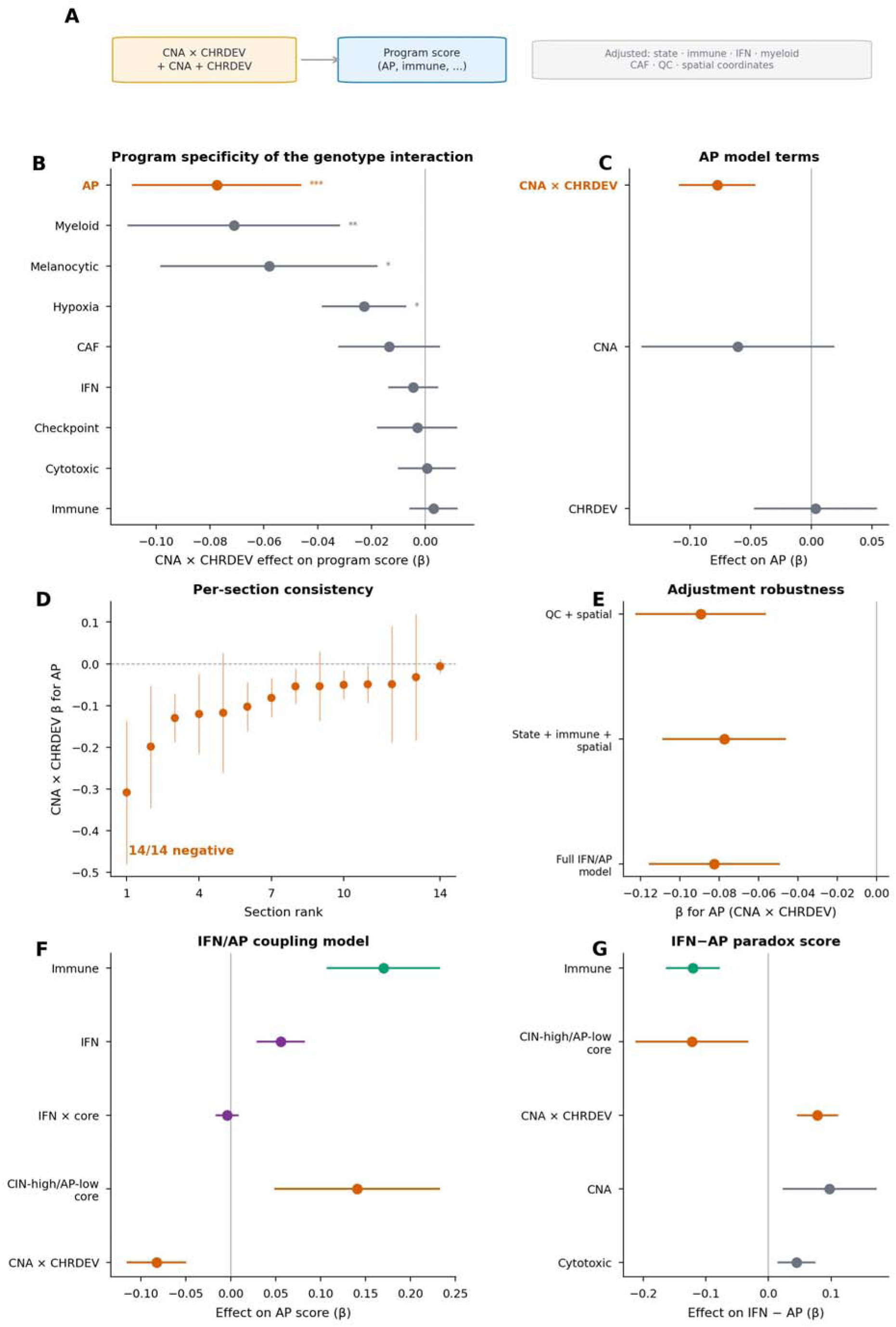
CNA-by-CHRDEV interaction is most specific for AP attenuation. a, Modeling scheme linking the CNA-by-CHRDEV interaction to AP score with covariate adjustment and section-level meta-analysis. b, Program-specific interaction effects across AP, immune, cytotoxic, checkpoint, IFN, CAF, hypoxia, melanocytic and myeloid programs. c, AP model terms for CHRDEV, CNA and CNA-by-CHRDEV. d, Section-level AP coefficients for the CNA-by-CHRDEV interaction. e, Covariate-adjustment robustness of the AP interaction. f, IFN/AP coupling model. g, IFN-minus-AP paradox model.

This interaction was more informative than either axis alone. CNA alone was not a stable negative AP predictor after adjustment, and chromosome-deviation alone captured broader stress and state variation. The interaction identified regions where copy-number-like expression imbalance and chromosome-scale expression deviation coincided. Across nine program outcomes, AP showed the most negative CNA-by-chromosome-deviation effect under the state-immune-spatial model, ranking first among AP, myeloid, melanocytic-state, hypoxia, CAF, IFN, checkpoint, cytotoxic and immune scores. This ranking supports the interpretation that the signal is not simply a generic consequence of low RNA quality or global program suppression. (Fig. 2b-e; Supplementary Fig. 4).

Several non-AP programs also showed associations with genotype imbalance, including myeloid and hypoxia-related features, consistent with the pleiotropic biology of chromosomal instability. The crucial observation is therefore not that AP was the only affected program, but that AP attenuation was directionally consistent, statistically robust and among the strongest program-level outcomes after extensive adjustment. This supports a mechanism in which transcriptome-inferred CIN-like fields are coupled to antigen-presentation erosion rather than merely representing a nonspecific stressed tissue compartment.

The adjusted modeling strategy was deliberately stricter than a simple AP-versus-CNA correlation. Models included local immune and IFN axes because these are expected biological drivers of AP expression; retaining the negative interaction after this adjustment means the genotype state is associated with lower AP beyond the AP-inducing inflammatory context. Models also included tumor-state and spatial-coordinate terms to reduce confounding by tumor-rich versus stromal-rich regions and by large-scale tissue gradients. Thus, the primary estimate should be interpreted as an adjusted within-section genotype-ecotype association, not as an uncorrected spot-level correlation.

### CIN-high/AP-low cores form AP-low neighborhoods that are depleted of immune and IFN signals

The next question was whether CIN-high/AP-low spots were isolated points or spatially organized tissue fields. Using k-nearest-neighbor topology within sections, CIN-high/AP-low cores were strongly enriched for AP-low neighbors. In the original topology analysis, AP-low neighbor enrichment across all melanoma sections had median z = 2.98 and Stouffer z = 18.8. In the final block-preserving spatial null across the 16 melanoma and breast sections, AP-low neighbor enrichment remained strong with Stouffer z = 26.66, median z = 4.99, p_high = 7.68 x 10^-157 and FDR = 9.21 x 10^-156. Leave-one-section, leave-one-dataset and leave-one-context analyses all preserved the expected direction. (Fig. 1g; Fig. 3b; Fig. 6c; Supplementary Fig. 6).

**Figure 3.**
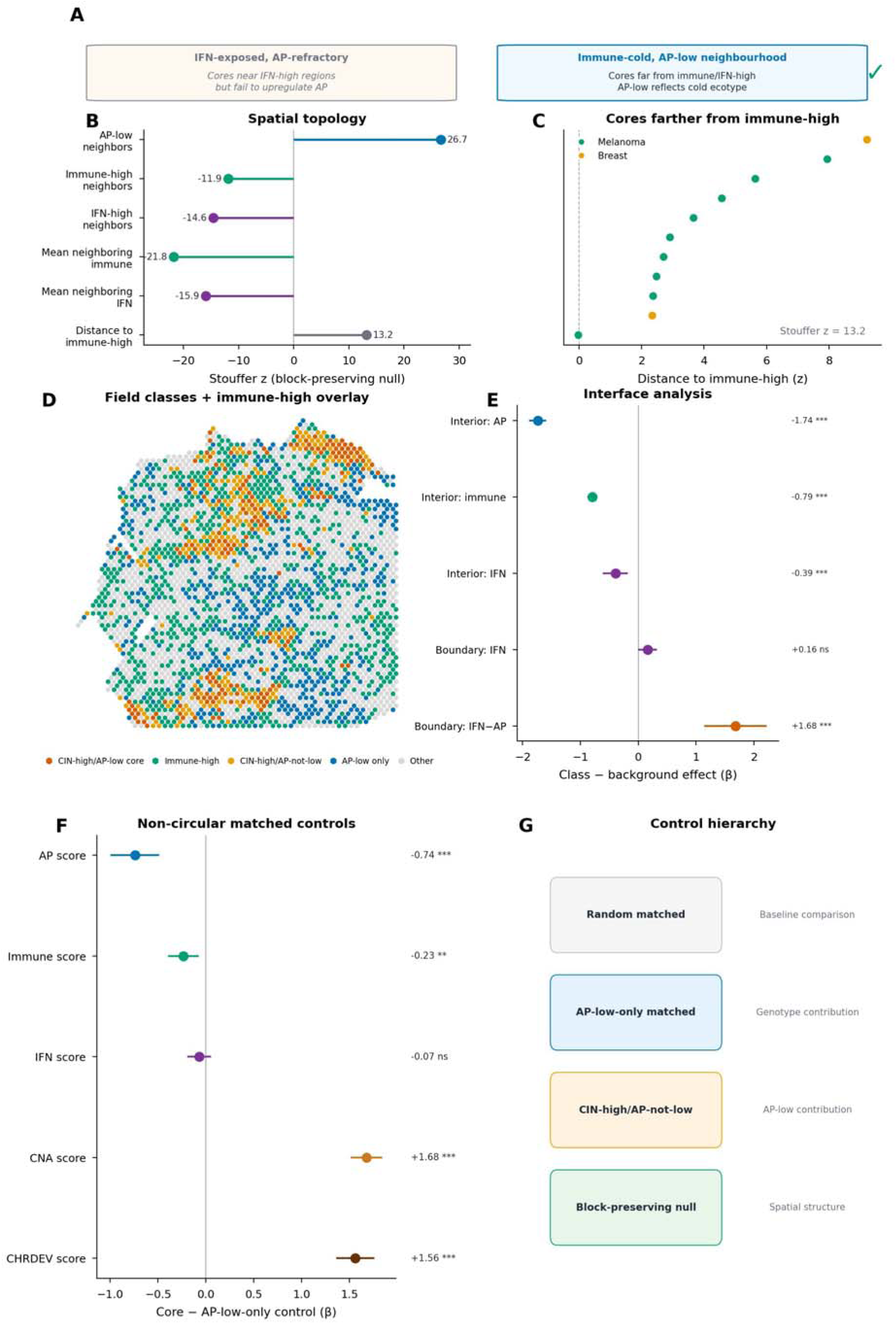
CIN-high/AP-low cores distinguish immune-cold interiors from IFN-exposed AP-low boundaries. a, Conceptual distinction between IFN-refractory AP-low boundaries and immune-cold AP-low interiors. b, Block-null topology around CIN-high/AP-low cores. c, Section-level distance-to-immune-high tests. d, Representative spatial interface map separating cores, immune-high neighborhoods and IFN-high neighborhoods. e, Interface contrasts for immune-cold AP-low interiors and IFN-high AP-low boundaries. f, Matched AP-low-only controls showing that cores remain distinct from other AP-low regions. g, Non-circularity control logic.

This AP-low topology alone could still be circular because AP-low is part of the core definition. Therefore, the analysis tested the non-defining immune and IFN neighborhoods around the same cores. Under block-preserving spatial nulls, CIN-high/AP-low cores were depleted of immune-high neighbors (Stouffer z = −11.87, FDR = 1.53 x 10^-32), depleted of IFN-high neighbors (Stouffer z = −14.57, FDR = 5.00 x 10^-48), had lower mean neighboring immune signal (Stouffer z = −21.78, FDR = 9.24 x 10^-105), and had lower mean neighboring IFN signal (Stouffer z = −15.93, FDR = 4.98 x 10^-57). They were also farther from immune-high regions than expected by the block null (Stouffer z = 13.21, FDR = 2.21 x 10^-39). (Fig. 3b,c).

These results reject the strongest version of the IFN-exposed AP-refractory interface model. In that model, CIN-high/AP-low regions should be located near IFN-high or immune-high neighborhoods while failing to raise AP. Instead, the dominant spatial pattern was immune-cold: CIN-high/AP-low cores were AP-low interiors with depleted immune and IFN neighborhood signals. A distinct IFN-high/AP-low boundary class existed, and its IFN-minus-AP score was elevated, but it did not explain the main CIN-high/AP-low topology. (Fig. 3a,d,e).

Ring-gradient analysis around CIN-high/AP-low cores made the field structure visually and quantitatively explicit. Across 16 sections, core-minus-far contrasts showed lower AP (beta = −2.192, FDR < 1 x 10^-300), lower immune signal (beta = −0.634, FDR < 1 x 10^-300) and lower IFN signal (beta = −0.393, FDR = 6.48 x 10^-5), while CNA and chromosome-deviation scores were higher in the cores (CNA beta = 2.468, FDR < 1 x 10^-300; chromosome-deviation beta = 1.846, FDR < 1 x 10^-300). The AP, CNA and chromosome-deviation gradients are partly expected from the core definition, but the immune and IFN gradients are not. These non-defining gradients provide the biologically informative evidence for immune-cold field organization. (Fig. 1h; Supplementary Fig. 5).

Boundary classification further separated immune-cold interiors from IFN-high AP-low interfaces. Immune-cold AP-low interiors had lower AP (beta = −1.739, FDR < 1 x 10^-300), lower immune signal (beta = −0.793, FDR < 1 x 10^-300) and lower IFN signal (beta = −0.394, FDR = 5.0 x 10^-4). In contrast, IFN-high/AP-low boundaries had an elevated IFN-minus-AP score (beta = 1.684, FDR = 2.3 x 10^-9). These results indicate that IFN-exposed AP-low boundaries exist, but the transcriptome-inferred CIN-high/AP-low field is dominated by immune-cold interiors rather than by IFN-rich boundaries. (Fig. 3d,e; Supplementary Fig. 5).

### Matched controls show that immune-cold topology is not only a consequence of AP-low selection

To test whether immune-cold topology merely reflected AP-low status, CIN-high/AP-low cores were compared with AP-low-only matched controls from the same sections. Controls were matched to preserve local context and available spot-level covariates while removing the requirement for simultaneous high CNA and high chromosome deviation. Relative to AP-low-only controls, CIN-high/AP-low cores had lower immune signal (beta = −0.2339, 95% CI −0.3936 to −0.0742, p = 0.0041, FDR = 0.00691, k = 16) and lower AP signal (beta = −0.7416, 95% CI −0.9951 to −0.488, FDR = 3.10 x 10^-8). IFN was directionally lower but not significant in this matched comparison (beta = −0.0707, 95% CI −0.193 to 0.052, p = 0.258, FDR = 0.302). (Fig. 3f; Fig. 6b).

A second control contrasted cores with spatially and technically matched random spots. CIN-high/AP-low cores again showed lower immune signal (beta = −0.2492, 95% CI −0.3566 to −0.1418, FDR = 1.28 x 10^-5) and lower AP signal (beta = −2.188, 95% CI −2.571 to −1.806, FDR = 2.10 x 10^-28). These matched-control results are important because they test features not fully used to define the cores. The AP component confirms the intended class identity, while the immune component shows that genotype-high AP-low cores are not interchangeable with all AP-low spots. (Fig. 6b).

The control analyses also expose a conservative limitation. Count-depth and detected-feature covariates differed in several matched contrasts, and therefore technical structure cannot be dismissed. For this reason, topology alone does not prove a cellular mechanism. Instead, spatial topology is interpreted together with adjusted regression, block nulls, leave-one stability, non-melanoma replication and malignant-cell single-cell analysis. (Fig. 6; Supplementary Fig. 6).

The matched-control results also clarify what should not be concluded. Because AP is part of the core definition, AP differences between cores and controls are expected and mainly verify class separation. By contrast, immune depletion relative to AP-low-only matched controls is not built into the class definition and is therefore more informative. The immune result shows that CIN-high/AP-low cores are not simply the lowest quartile of AP redistributed across tissue; they are a subset of AP-low regions with additional genotype-high and immune-cold features.

The block nulls and matched controls were complementary: the former tested whether core neighborhoods were more AP-low or immune-cold than expected under coarse tissue geography, whereas the latter tested whether cores differed from plausible alternative spot classes within the same section. Their agreement supports a non-random genotype-high AP-low region with non-defining immune-cold features.

The IFN result was more heterogeneous than the immune result: IFN depletion was directionally consistent but not always significant in AP-low-only matched contrasts, indicating that low immune neighborhood exposure is the more stable non-defining property of the cores.

### Breast cancer Visium sections reproduce AP-low immune-cold topology but not a fully significant interaction

To address melanoma specificity, the spatial framework was applied to two public 10x Genomics breast cancer Visium sections. These data were not treated as a pan-cancer proof, because two sections cannot establish universality. They were used as a concrete non-melanoma test of whether the same spatial topology can emerge in another epithelial cancer context. The strongest conserved signal was AP-low neighborhood topology: breast CIN-high/AP-low cores showed median z = 23.69 and Stouffer z = 31.4 for AP-low neighbor enrichment, with p_high < 1 x 10^-200 and FDR < 1 x 10^-200 across the two sections. (Fig. 4a-d; Supplementary Fig. 3).

**Figure 4.**
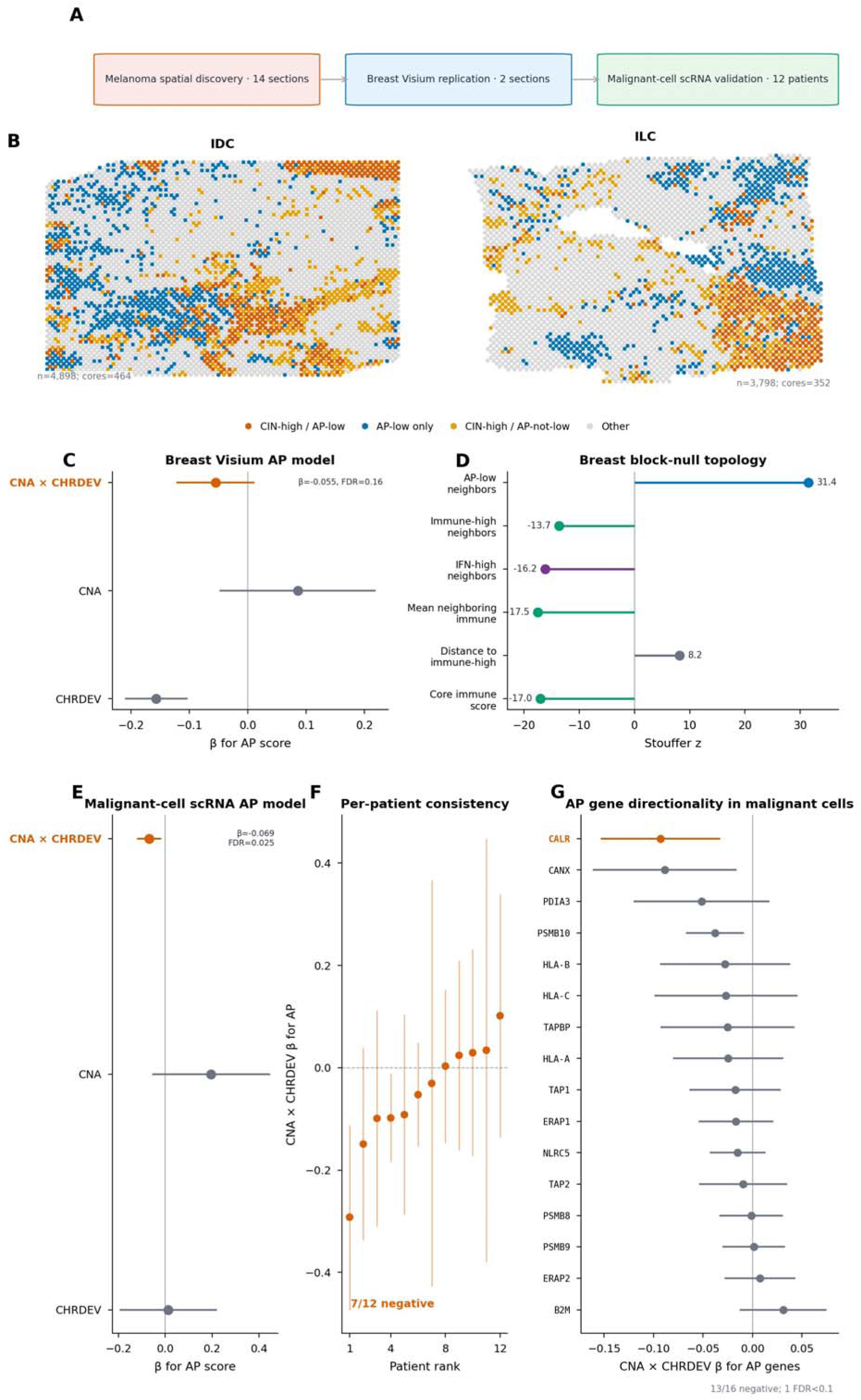
Cross-context validation in breast spatial data and malignant-cell scRNA-seq. a, Validation design linking melanoma spatial discovery to breast spatial sections and malignant-cell melanoma scRNA-seq. b, Breast IDC and ILC spatial field maps. c, Breast spatial AP model terms. d, Breast spatial topology showing AP-low neighbor enrichment and immune/IFN depletion. e, Malignant-cell scRNA-seq AP model terms. f, Patient-level CNA-by-CHRDEV coefficients for AP in malignant cells. g, AP gene-level CNA-by-CHRDEV coefficients in malignant cells.

The breast sections also showed immune-cold topology. Neighbor immune-high fraction was depleted (Stouffer z = −13.66), and the final block-null analysis across contexts showed strong depletion of immune and IFN neighborhood means. Matched-control contrasts in breast were directionally consistent: relative to AP-low-only matched controls, cores had lower IFN and immune signal, although the small number of sections limited inference. The non-melanoma spatial result therefore supports conservation of the spatial neighborhood phenotype rather than definitive conservation of every regression coefficient.

The genotype interaction itself was weaker in breast. The non-melanoma CNA-by-chromosome-deviation term for AP was negative in both sections but did not reach significance by random-effects meta-analysis (beta = −0.0550, 95% CI −0.122 to 0.0121, p = 0.108, FDR = 0.162). In contrast, chromosome deviation alone was strongly negative (beta = −0.1568, 95% CI −0.2105 to −0.1032, p = 9.88 x 10^-9, FDR = 2.96 x 10^-8). The appropriate conclusion is therefore partial cross-cancer conservation: breast sections reproduce AP-low immune-cold topology, while the continuous genotype interaction requires larger non-melanoma cohorts. (Fig. 4c,d).

The breast sections were most useful as an independent topology test. They do not establish that melanoma and breast tumors share identical mechanisms, but they show that the central spatial endpoint - AP-low neighborhoods depleted of immune and IFN signal - can be recovered outside melanoma despite a weaker continuous interaction.

### Malignant-cell single-cell RNA-seq supports a tumor-intrinsic genotype-conditioned AP association

A major alternative explanation for spot-level spatial coupling is spot mixture. CNA-like signal may be driven by malignant cells, whereas AP or immune signal could arise from non-malignant cells in the same spot. To address this, malignant-cell single-cell RNA-seq from metastatic melanoma was analyzed independently. Cells were stratified by patient, malignant identity and inferred genotype-state features, and AP activity was modeled as a function of CNA, chromosome deviation and the CNA-by-chromosome-deviation interaction with available covariates.

Across 12 patient-level malignant-cell models, the CNA-by-chromosome-deviation interaction was negatively associated with AP activity (beta = −0.0685, 95% CI −0.1194 to −0.0177, p = 0.00825, FDR = 0.0247). CNA alone was not negative and was directionally positive in the meta-analysis (beta = 0.1956, p = 0.128), emphasizing again that the relevant signal is not a simple monotonic CNA/AP relationship. The interaction result supports the idea that AP attenuation occurs within malignant-cell genotype-state space and is not solely an artifact of mixed spatial spots. (Fig. 4e,f; Supplementary Fig. 7).

Gene-level AP coherence was more nuanced. In malignant-cell single-cell data, 13 of 16 AP machinery genes were directionally negative for the CNA-by-chromosome-deviation term, but only one reached FDR < 0.1. In spatial spot data, individual AP genes did not show coherent negative gene-level effects; the CNA-by-CHRDEV coefficients for individual AP genes were predominantly positive at the spot level (Supplementary Fig. 7d), likely reflecting confounding by spot composition and immune-cell AP expression that the pathway-level score and covariate adjustment can absorb but individual gene models cannot. Therefore, the main claim remains pathway-level AP attenuation rather than collapse of each AP machinery gene. This distinction is important: the discovery concerns a coordinated program score and tissue architecture, not uniform repression of every antigen-processing component. (Fig. 4g; Supplementary Fig. 7).

The malignant-cell validation also provided a useful negative control on overstatement. If CNA alone were sufficient to explain AP attenuation, a negative CNA main effect would be expected in malignant cells. Instead, the CNA main effect was not negative, whereas the interaction with chromosome deviation was negative. This mirrors the spatial finding and supports the interpretation that AP attenuation is linked to a combined genotype-state imbalance rather than to copy-number burden in isolation.

### Patient-level outcome analyses support clinical relevance without defining a biomarker

The spatial discovery was projected onto patient-level TCGA cohorts to ask whether the genotype-ecotype state has prognostic relevance. In TCGA-SKCM, a bulk adverse score combining high CNA, high chromosome deviation and low AP was associated with worse overall survival after adjustment for age and stage (HR = 1.46, 95% CI 1.25-1.71, p = 2.7 x 10^-6, FDR = 6.91 x 10^-5; n = 320, 168 events). The result was similar with additional TMB adjustment (HR = 1.45, 95% CI 1.24-1.70, p = 4.0 x 10^-6). AP-low and chromosome-deviation features individually also carried prognostic signal in SKCM. (Fig. 5a-c; Supplementary Fig. 8).

**Figure 5.**
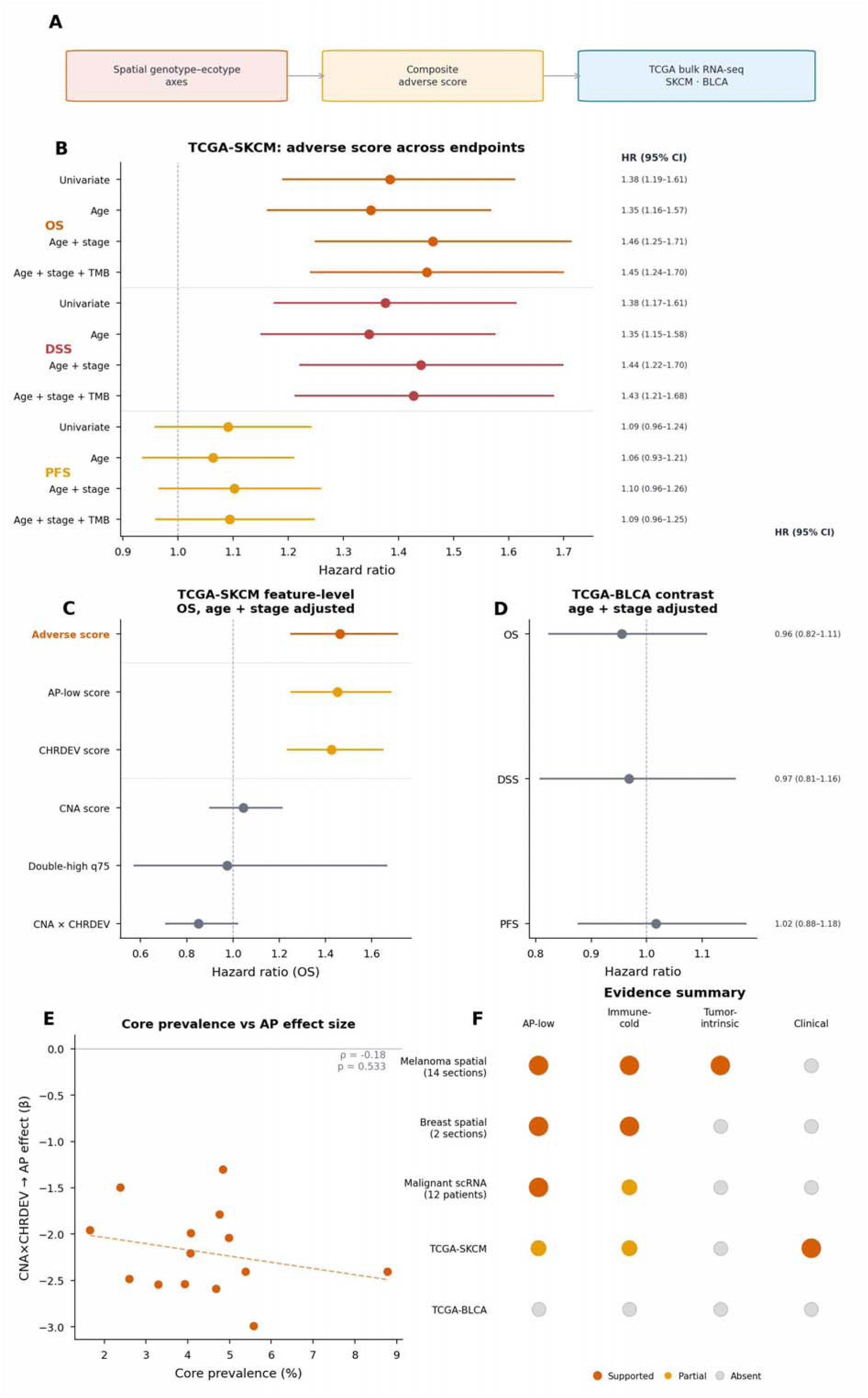
Clinical projection of the spatial genotype-ecotype state. a, Projection of spatial genotype-ecotype axes into TCGA bulk RNA-seq cohorts. b, TCGA-SKCM overall-survival hazard ratios for the adverse genotype-ecotype score across adjustment sets. c, TCGA-SKCM feature-level survival models. d, TCGA-BLCA survival projection. e, Section-level association between core prevalence and CNA-by-CHRDEV AP effect size. f, Evidence matrix summarizing spatial discovery, breast spatial replication, malignant-cell validation and clinical projection.

**Figure 6.**
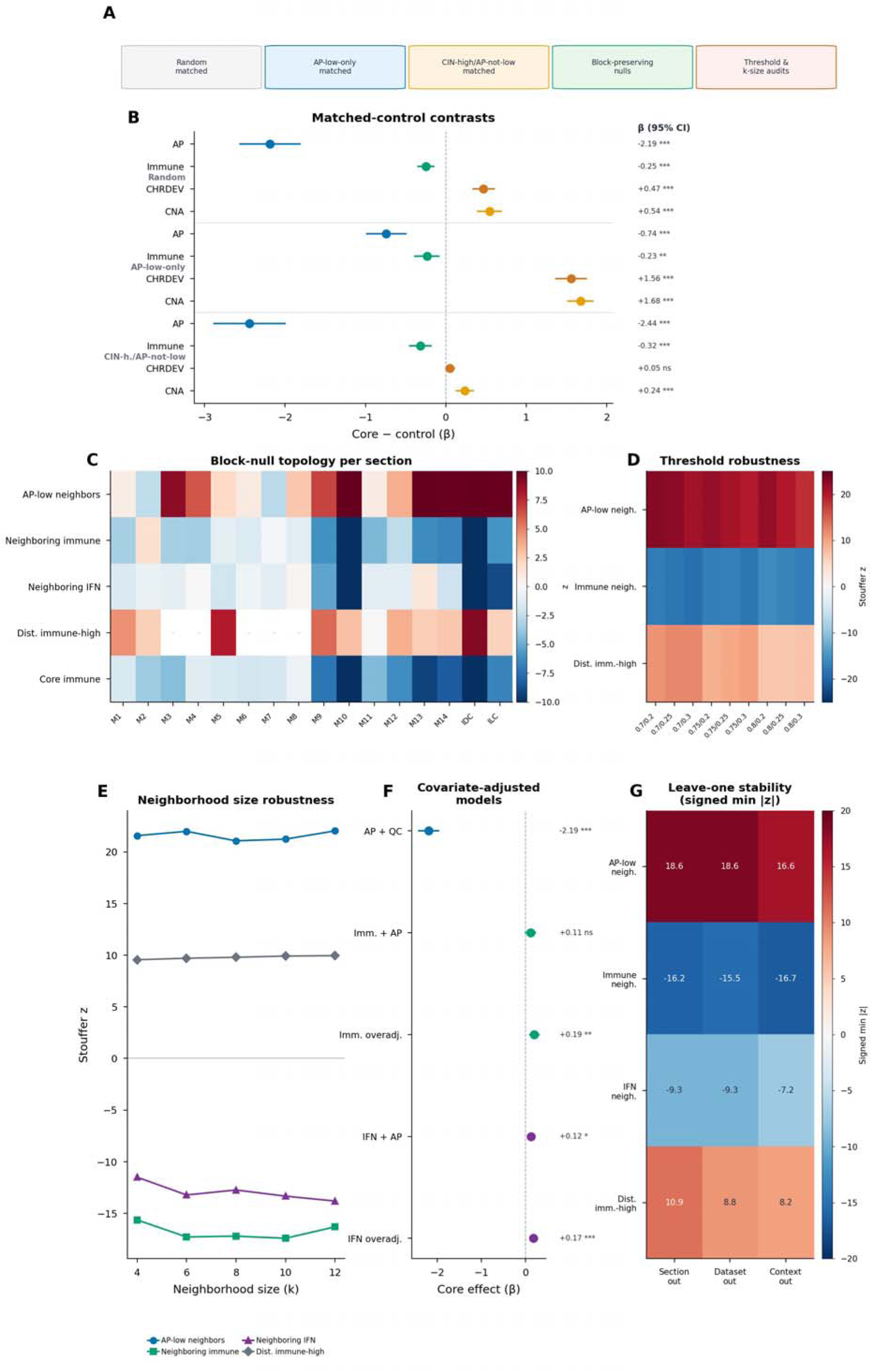
Robustness and non-circularity controls. a, Control framework for matched random, AP-low-only, CIN-high/AP-not-low and block-spatial null analyses. b, Matched-control contrasts across programs. c, Per-section block-null topology heatmap. d, Threshold robustness across alternative high-CNA/high-CHRDEV and AP-low definitions. e, Neighborhood-size robustness across k-nearest-neighbor definitions. f, Covariate-adjusted models testing CIN-high/AP-low core effects on AP and immune/IFN depletion under progressively strict adjustment sets, including over-adjusted models that control for the outcome variable itself as a stress test. g, Leave-one-section, leave-one-dataset and leave-one-context stability.

TCGA-BLCA did not show the same adverse-score association with overall survival (HR = 0.955, p = 0.549), despite earlier bulk evidence of continuous AP/genotype coupling. This discordance reinforces the context-dependent interpretation. The spatial architecture has strong melanoma evidence, breast topology support and single-cell tumor-intrinsic support, but patient-level clinical consequences vary by disease context and endpoint. (Fig. 5d).

The melanoma immunotherapy projection remains supportive rather than definitive. Previously analyzed ICI response data showed a directional adverse-score signal (AUC = 0.627, logistic p = 0.0072, FDR = 0.072 after multiple-testing correction across prespecified tests), below the threshold for a predictive biomarker. Accordingly, clinical analyses are presented as patient-level projection of the spatial biology, not as a validated ICI response classifier. (Supplementary Fig. 9).

Survival modeling was intentionally separated from the spatial discovery. Spatial topology cannot be reconstructed from bulk TCGA samples. Instead, the TCGA analysis asks whether a bulk projection of the same adverse genotype-ecotype axis has patient-level consequences. The positive TCGA-SKCM result strengthens biological relevance, whereas the negative BLCA result prevents overgeneralization. This balance is important for the biology: clinical projection can support relevance, but it should not replace direct tissue-level evidence. (Fig. 5f; Supplementary Fig. 10).

This final layer places the spatial state in a patient-level context. The survival association in melanoma indicates that the same genotype-ecotype axis can be detected in bulk cohorts with clinical outcomes, while the absence of a matching BLCA survival signal shows that downstream consequences are disease-context dependent.

## Discussion

This study supports a biological model in which transcriptome-inferred CIN-like genotype fields mark AP-low, immune-cold spatial neighborhoods. The model differs from a bulk aneuploidy-immune-cold correlation in three ways. First, the unit of analysis is a spatial field within tissue sections. Second, the relevant genotype state is the combination of inferred CNA-like burden and chromosome-expression deviation rather than CNA alone. Third, the dominant topology is not IFN-rich AP-refractory boundary behavior but immune-cold AP-low neighborhood organization.

The main conceptual advance is the field-level framing. To our knowledge, no prior study has systematically tested whether expression-inferred CNA states predict spot-level AP attenuation within spatial neighborhoods using block-preserving nulls and matched controls. Prior studies linked aneuploidy and immune evasion at the bulk tumor level and established antigen-presentation loss as a mechanism of immune escape. The present analysis adds spatial organization: CNA/CHRDEV-high regions are not only AP-low but are embedded in AP-low immune-cold neighborhoods. This difference matters because immune escape in tissue is not just a molecular state; it is also a geography that determines where immune cells encounter tumor cells and where inflammatory signals are absent or ineffective.

The strongest evidence comes from the convergence of independent tests. The melanoma spatial interaction was negative for AP in all 14 sections after state, immune, QC and coordinate adjustment. CIN-high/AP-low cores were enriched for AP-low neighbors under block-preserving spatial nulls and depleted of immune and IFN neighbors. Matched controls showed that immune depletion was not simply a property of all AP-low spots. Breast cancer sections reproduced AP-low immune-cold topology, and malignant-cell single-cell RNA-seq supported a tumor-intrinsic CNA-by-chromosome-deviation association with AP repression. TCGA-SKCM survival analyses linked the adverse genotype-ecotype score to worse patient outcome. No single layer proves the mechanism, but together they make a coherent spatial-genomic argument.

The findings refine how chromosomal-instability-like genotype states may shape immune escape when measured from spatial transcriptomes. Chromosomal instability can influence tumor biology through gene dosage, stress signaling, altered cell state, metastasis-associated programs and immune interactions 5–9. The present analysis suggests that expression-derived proxies of these processes can become geographically organized in tissue. A high-CNA/high-chromosome-deviation state does not merely associate with lower AP in isolation; it marks regions in which AP-low malignant-state programs and low immune/IFN neighborhood exposure co-localize. This provides a tissue-level explanation for why bulk aneuploidy may correlate with low immune cytolytic activity: part of the signal may arise from spatially expanded immune-cold AP-low fields.

The non-melanoma breast validation should be read as topology replication rather than as a completed cross-cancer atlas. Two independent breast sections reproduced AP-low immune-cold neighborhood organization, but the continuous interaction requires larger non-melanoma cohorts. The result therefore supports partial conservation while keeping the strongest claim anchored in melanoma.

The results also clarify the IFN/AP uncoupling question. Although IFN-high AP-low interfaces were detectable as a separate class, they did not account for the dominant CIN-high/AP-low genotype field. Instead, the core regions were depleted of IFN-high neighbors and farther from immune-high neighborhoods, supporting immune-cold topology rather than generalized IFN-refractory AP failure.

The single-cell result addresses the key spot-mixing criticism. Malignant-cell scRNA-seq cannot make Visium single-cell resolved, but it shows that the genotype-conditioned AP association is detectable within malignant cells, supporting a tumor-intrinsic component to the spatial pattern.

Several limitations should constrain interpretation. Visium spots are mixtures, and the CNA and CHRDEV axes are transcriptome-derived proxies rather than DNA-based clone calls or direct live-CIN measurements. The non-melanoma spatial validation currently includes only two breast sections, so it establishes plausibility and topology conservation but not pan-cancer generality. AP gene-level coherence was not strong in spatial data, and clinical projections are not predictive biomarkers. The study therefore makes a precise claim: transcriptome-inferred CNA/CHRDEV-high fields are associated with pathway-level AP attenuation and immune-cold spatial neighborhoods, with strongest evidence in melanoma and partial conservation in breast spatial data.

Future work should test the model with spatial multi-omics that measures copy number, transcriptome, protein-level HLA expression and immune cell phenotypes in the same tissue sections. DNA-based clone maps could determine whether the inferred genotype fields correspond to true subclonal expansions. Multiplexed imaging could resolve whether AP-low cores contain tumor cells with reduced HLA protein or whether AP signal is diluted by local composition. Functional experiments could test whether induced CIN or selected aneuploid states actively remodel immune neighborhoods. A 50-gene malignant-cell signature upregulated in CIN-high/AP-low core cells (Supplementary Fig. 10c) may provide a starting point for targeted validation of the core state in independent cohorts. The present study provides a computationally rigorous map of the phenomenon and a framework for those mechanistic experiments.

Translationally, the field model suggests that AP restoration and immune recruitment may need to be considered together. Restoring AP in isolated tumor cells may not correct immune-cold genotype-high neighborhoods, while immune recruitment alone may be insufficient if genotype-high malignant cells have reduced AP inducibility. These remain hypotheses, because the present study is observational.

Finally, chromosome-scale expression deviation may capture biology not reducible to canonical CNA burden. The strongest signals repeatedly involved the CNA-by-chromosome-deviation interaction, suggesting that immune topology emerges from a combined copy-number-like and transcriptional-instability state. Direct DNA copy-number maps, allele-specific HLA calls and matched protein imaging will be needed to identify causal components.

## Methods

### Spatial datasets and preprocessing

Public melanoma and breast cancer spatial transcriptomic datasets were processed as spot-level count matrices with associated tissue coordinates. The melanoma discovery set contained 29,341 spots from 14 tissue sections across three GEO datasets: GSE250636 (GSM7983358_sample2, GSM7983359_sample4, GSM7983360_sample7, GSM7983361_sample8, GSM7983362_sample12, GSM7983363_sample13, GSM7983364_sample14, GSM7983365_sample15 and GSM7983366_sample16), GSE271422 (GSM8376639_sample1, GSM8376640_sample9 and GSM8376641_sample10) and GSE316760 (GSM9459774_mel2 and GSM9459775_mel3). Non-melanoma spatial validation used two 10x Genomics breast cancer Visium sections comprising 8,696 spots: Human Breast Cancer, Ductal Carcinoma In Situ/Invasive Carcinoma, FFPE, and Human Breast Cancer, Block A Section 1. Coordinates were standardized within sections. Low-information spots were removed using available in-tissue, count-depth and detected-feature metadata. Gene symbols were harmonized to a common human gene-symbol namespace, and module scores were computed from log-normalized expression where raw count matrices were available. Probabilistic spot deconvolution (e.g., Cell2location, RCTD) was not applied because the analysis framework uses continuous module scores with immune, IFN and tumor-state covariates rather than categorical cell-type fractions, and because the single-cell validation provides an independent tumor-intrinsic control for spot-mixture confounding.

### Program scoring

Antigen-presentation activity was summarized from MHC-I processing and presentation genes including HLA-A, HLA-B, HLA-C, B2M, TAP1, TAP2, TAPBP, PSMB8, PSMB9, PSMB10, ERAP1, ERAP2, NLRC5, CALR, CANX and PDIA3. For a gene set G and spot i in section s, the module score was computed as the mean of available log-expression values for genes in G and then standardized within section as z_s(x_i) = (x_i - mean_s(x))/sd_s(x). Immune, cytotoxic, IFN, myeloid, CAF, endothelial, hypoxia, checkpoint and tumor-state scores were computed from curated gene sets using the same section-wise standardization. The AP score was treated as a pathway-level activity measure. Individual AP gene analyses were performed separately and were not used to support the main pathway claim unless multiple genes showed coherent effects.

### Genotype-state proxies

The CNA axis was an inferred copy-number-like expression-shift score selected from the available Stage31 columns, prioritizing infercnv_like_shift_z_no_chr6_se7_immune or its cna_chr_imbalance equivalent when present. This choice excluded chromosome 6/MHC and AP/immune leakage where possible, reducing circularity between the genotype proxy and the AP outcome. Chromosome-level expression shifts were computed for autosomes with sufficient genes by averaging standardized expression within each chromosome. For spot i and chromosome c, E_ic denotes the chromosome-level expression-shift score. The primary chromosome-deviation score was CHRDEV_i = z_s(mean_c |E_ic|), with max, top-three and variance summaries retained as sensitivity features. Both CNA and CHRDEV were z-scored within tissue section or patient, using the section mean as the reference rather than normal-tissue spots (which were not available in all datasets). The interaction term was CNA_z x CHRDEV_z. CIN-high/AP-low cores were defined within each section as spots with CNA_z and CHRDEV_z above their section-specific 75th percentiles and AP_z below its 25th percentile. This is a transcriptome-inferred CIN-like state, not a direct DNA copy-number or live-CIN measurement.

### Adjusted spatial regression

For each section, AP_z was modeled as AP_z = beta0 + beta1*CNA_z + beta2*CHRDEV_z + beta3*(CNA_z x CHRDEV_z) + betaC*C + epsilon, where C contained available immune, IFN, cytotoxic, myeloid, CAF, hypoxia, checkpoint and tumor-state scores, count-depth and detected-feature covariates, and spatial coordinate basis terms. Linear models were fit section by section using ordinary least squares with HC3 robust standard errors. The primary coefficient was beta3 for the CNA-by-CHRDEV interaction. Section-level estimates were combined using random-effects meta-analysis. Multiple testing was controlled with the Benjamini-Hochberg false discovery rate procedure. Reported FDR values of zero from numerical underflow were replaced with conservative bounds such as FDR < 1 x 10^-300.

### Spatial topology and null models

Neighborhoods were defined using k-nearest-neighbor graphs within each tissue section; the default rigor audit used k = 6 nearest neighbors, with neighborhood-size sensitivity across k = 4, 6, 8, 10 and 12. For each CIN-high/AP-low core set, the analysis computed neighbor AP-low fraction, neighbor immune-high fraction, neighbor IFN-high fraction, mean neighboring immune and IFN signals, core mean immune and IFN signals, and mean distance to immune-high spots. Block-preserving nulls divided each section into a 4 x 4 grid using quantile bins of standardized x and y coordinates. For each permutation, the number of core spots per spatial block was preserved and matched null spots were drawn from non-core spots in the same block, with replacement only when the block-specific pool was smaller than required. The final Stage47 analysis used 500 permutations and required at least five core spots per section. Section-level observed-minus-null z-scores were combined by Stouffer’s method with directional p values.

### Ring-gradient and interface analyses

For each section, CIN-high/AP-low core spots were used as anchors. Neighbor rings were generated from graph distance from these anchors and summarized as core, ring 1, ring 2, ring 3 and far-field regions. Program means were computed per ring and per section, and core-minus-far contrasts were combined by random-effects meta-analysis. Interface classes were defined by combinations of AP-low status, immune/IFN neighborhood scores and CIN-high genotype status. The principal classes were immune-cold AP-low interiors, IFN-high AP-low boundaries, immune-high boundaries and background. These classes were used to distinguish immune-cold interiors from IFN-high/AP-low boundaries without assuming that all AP-low regions share the same inflammatory context.

### Matched-control analyses

To test non-circularity, each CIN-high/AP-low core was compared with matched control spots from the same section. AP-low-only controls were AP-low but not CIN-high, while CIN-high/AP-not-low controls were CNA/CHRDEV-high but not AP-low. Random matched controls were sampled from the remaining section background. Matching used nearest-neighbor matching in the standardized covariate space defined by x coordinate, y coordinate, tumor-state score, count-depth score and detected-feature score when available. Missing values were replaced by section medians, and matching variables were z-standardized before nearest-neighbor search. Outcomes included AP, immune, IFN, CNA, chromosome deviation, tumor-state and QC scores. These paired core-minus-control contrasts were summarized per section and meta-analyzed across sections and contexts.

### AP gene-level analysis

Individual antigen-presentation genes were tested separately in spatial and malignant-cell datasets when expression values were available. Per-gene models used the same CNA, chromosome-deviation and interaction structure as the pathway-level models. Results were interpreted conservatively: individual-gene coherence was considered supportive only if several genes showed negative interaction effects after FDR correction. Because this condition was not met in spatial data, the interpretation remains at the pathway-level AP scale.

### Single-cell validation

Malignant-cell single-cell RNA-seq from metastatic melanoma was analyzed independently using the processed GSE72056/Tirosh matrix. Malignant cells were identified from processed metadata when available; if malignant metadata were unavailable or insufficient, all melanoma tumor-cell matrix columns passing the expression filters were retained and the uncertainty was recorded in the inventory table. Cells were assigned to patients from cell-name prefixes. For expression-derived copy-number-like axes, chromosome-level expression shifts were computed relative to the non-malignant reference cells when at least 100 reference cells were available, otherwise relative to the dataset mean. CNA_expr_burden was the mean absolute chromosome shift across autosomes, and CHRDEV_top3_axis was the mean of the three largest absolute chromosome shifts. AP activity, IFN score, cell-cycle score, immune score and detected-gene burden were standardized within patient. AP_z was then modeled per patient as a function of CNA_z, CHRDEV_z, CNA_z x CHRDEV_z, IFN_z, cell_cycle_z, immune_z and detected-gene z-score using HC3 robust standard errors. Patient-level coefficients were combined by random-effects meta-analysis. Gene-level AP analyses were conducted for individual antigen-processing genes as a secondary test.

### Clinical projection

TCGA-SKCM and TCGA-BLCA feature tables were linked to cBioPortal PanCancer Atlas clinical outcome data. Overall survival, disease-specific survival and progression-free survival endpoints were harmonized from available clinical fields. Cox proportional hazards models tested standardized adverse genotype-ecotype scores and component features with adjustment for age, stage and TMB where available. The adverse score was a bulk projection of the spatial genotype-ecotype state and was not used to infer spatial topology in TCGA. ICI response analyses were treated as projection tests using available public melanoma cohorts and were not optimized as predictive classifiers.

### Reporting and reproducibility

All analyses were performed with fixed thresholds defined before interpretation. Results are reported as section-level or patient-level meta-analytic estimates rather than only spot-level p values. Interpretations were assigned according to evidence strength: primary spatial discovery in melanoma, partial non-melanoma spatial conservation, orthogonal malignant-cell support and patient-level prognostic projection. The associated figure package contains vector figure files, generation scripts for the six main figures and a data-to-figure map. The source-data transfer archive contains the Stage31, Stage38, Stage41C, Stage42, Stage43C, Stage44/45, Stage46, Stage47 and Stage48 scripts, logs and result tables used for the reported analyses.

### Quality-control and sensitivity summaries

Analyses emphasized section-level replication, leave-one stability and multiple null formulations. Leave-one-section, leave-one-dataset and leave-one-context summaries were computed for matched controls and block-null topology. Effects were promoted to main-text claims only when they showed the expected direction, were statistically supported after multiple-testing correction and were stable to leave-one analyses. Directional or underpowered results were reported as supportive or context-dependent rather than as primary discoveries.

### Inference criteria

Primary spatial evidence required concordance of adjusted melanoma interaction models, topology under block-preserving nulls, non-circular matched-control behavior and leave-one stability. Non-melanoma breast data were interpreted as supportive when AP-low immune-cold topology was reproduced, even if the continuous interaction remained underpowered. Single-cell RNA-seq was interpreted as tumor-intrinsic support when malignant-cell AP decreased with the CNA-by-chromosome-deviation interaction. Clinical analyses were interpreted only as patient-level projection and were not used to define the spatial mechanism.

### Output reproducibility

Figure-ready summaries were generated from the same section-level and patient-level tables used for statistical inference. Full model inventories, AP gene-level tests, leave-one stability summaries and dataset-specific coefficients are provided as Supplementary Tables 1–13 so that the main visualizations can be traced back to numerical outputs. Figures 1-6 and Supplementary Figs. 1-10 are mapped to their source data in the Supplementary Information.

## Data availability

The spatial transcriptomic, single-cell RNA-seq, TCGA and cBioPortal datasets analyzed in this study are publicly available from the original repositories and study accessions cited in the References. The melanoma spatial discovery set used GEO accessions GSE250636, GSE271422 and GSE316760 with the exact section labels listed in Methods. Breast validation used the 10x Genomics Visium Human Breast Cancer DCIS/Invasive Carcinoma FFPE and Human Breast Cancer Block A Section 1 datasets. The malignant-cell single-cell analysis used GSE72056. TCGA clinical data were accessed through cBioPortal PanCancer Atlas resources. Processed source tables, harmonized program scores, section-level model outputs, figure-ready summaries and figure-generation scripts are included in the associated source-data/figure package for preprint review; a permanent public repository is being prepared for journal submission.

## Code availability

Analysis scripts used to generate the final tables and figures are included in the associated source-data transfer archive and figure package. The code records fixed thresholds, model formulas, covariate sets, null-model settings, permutation counts and random seeds used for the reported analyses. A permanent GitHub or Zenodo repository is being prepared for journal submission.

## Acknowledgements

The author thanks the investigators who generated the public spatial transcriptomic, single-cell and TCGA resources analyzed here. For this work the HPC-cluster Hummel-2 at University of Hamburg was used. The cluster was funded by Deutsche Forschungsgemeinschaft (DFG, German Research Foundation) – 498394658. No new human subject data were generated for this study.

## Author contributions

S.D. conceived the study, designed the analyses, performed the computational work, interpreted the results and wrote the manuscript.

## Supplementary Figures

**Supplementary Figure 1:**
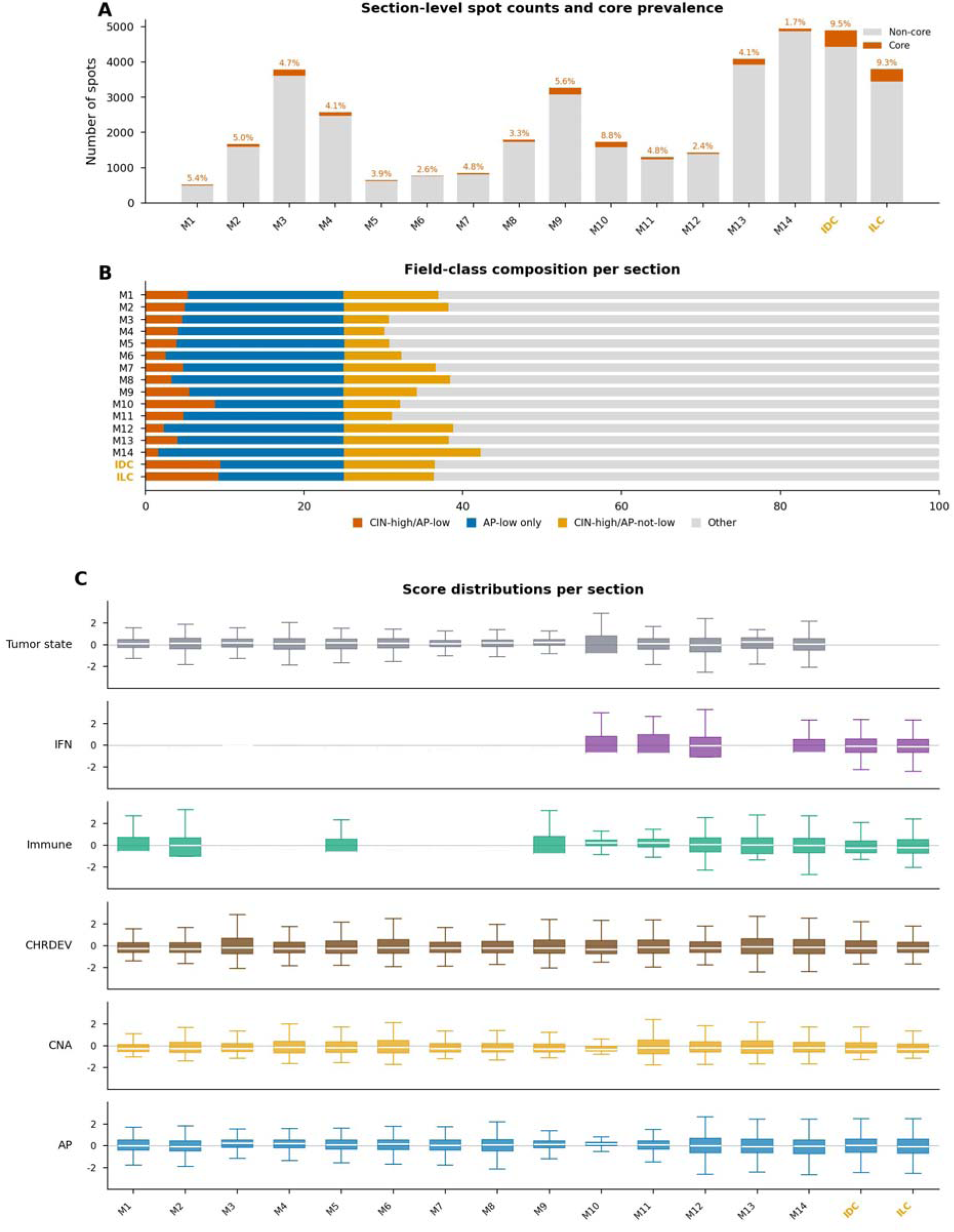
Dataset inventory, field-class composition, and score distributions Section-level spot counts and core prevalence

**Supplementary Figure 2:**
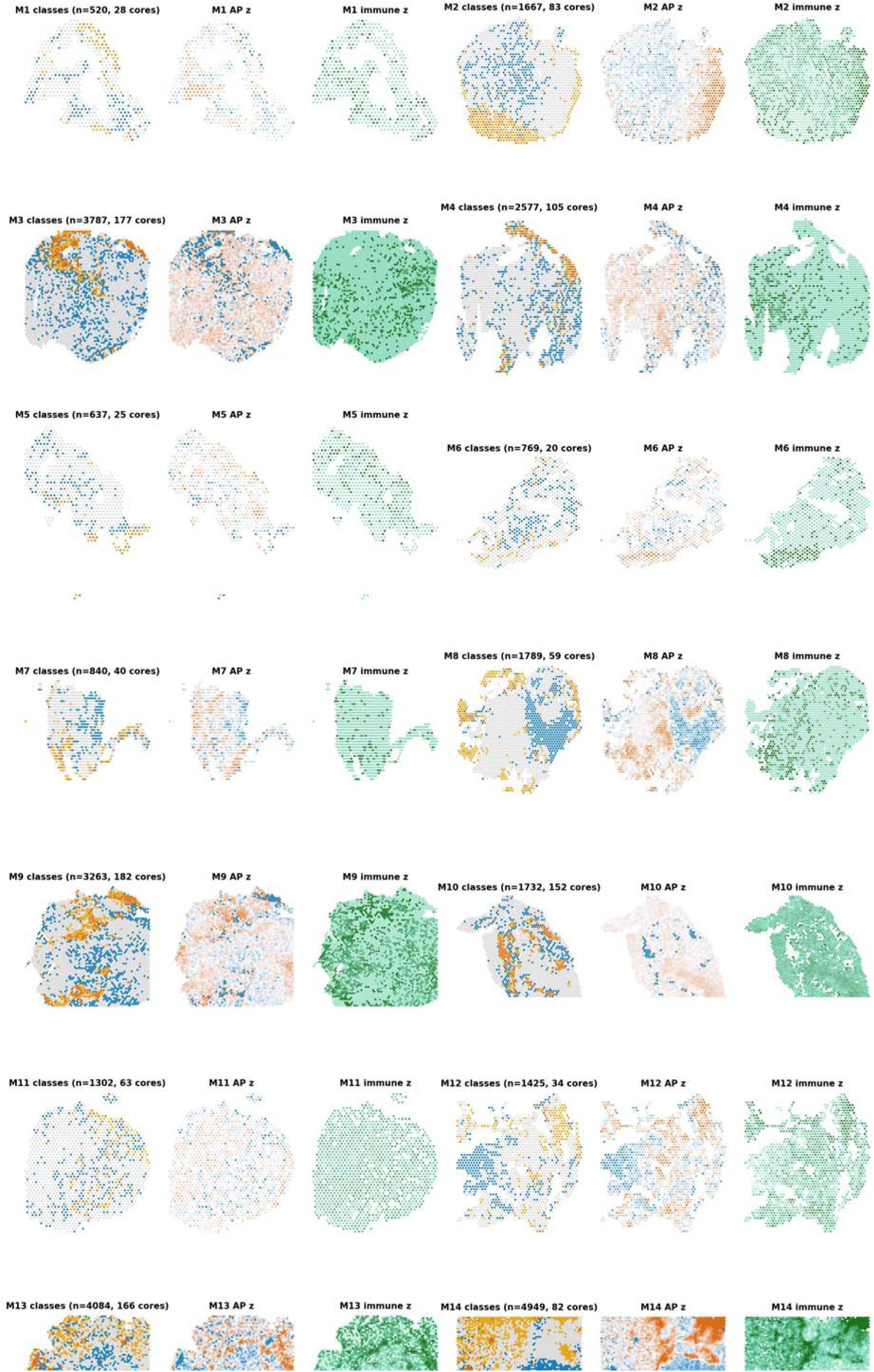
All melanoma spatial sections - field classes, AP z-score, immune z-score

**Supplementary Figure 3:**
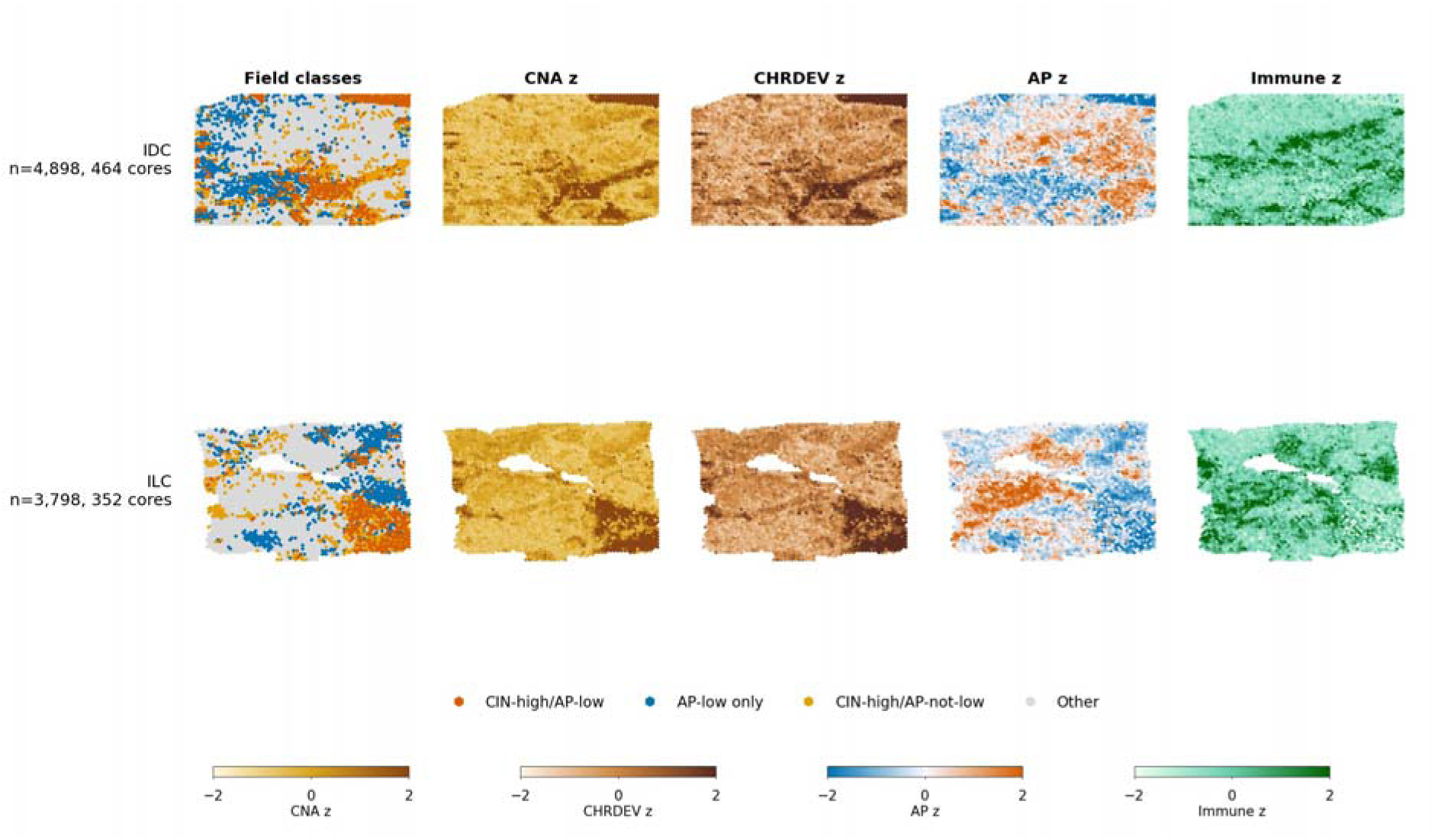
Breast spatial replication - field classes, CNA, CHRDEV, AP, immune

**Supplementary Figure 4:**
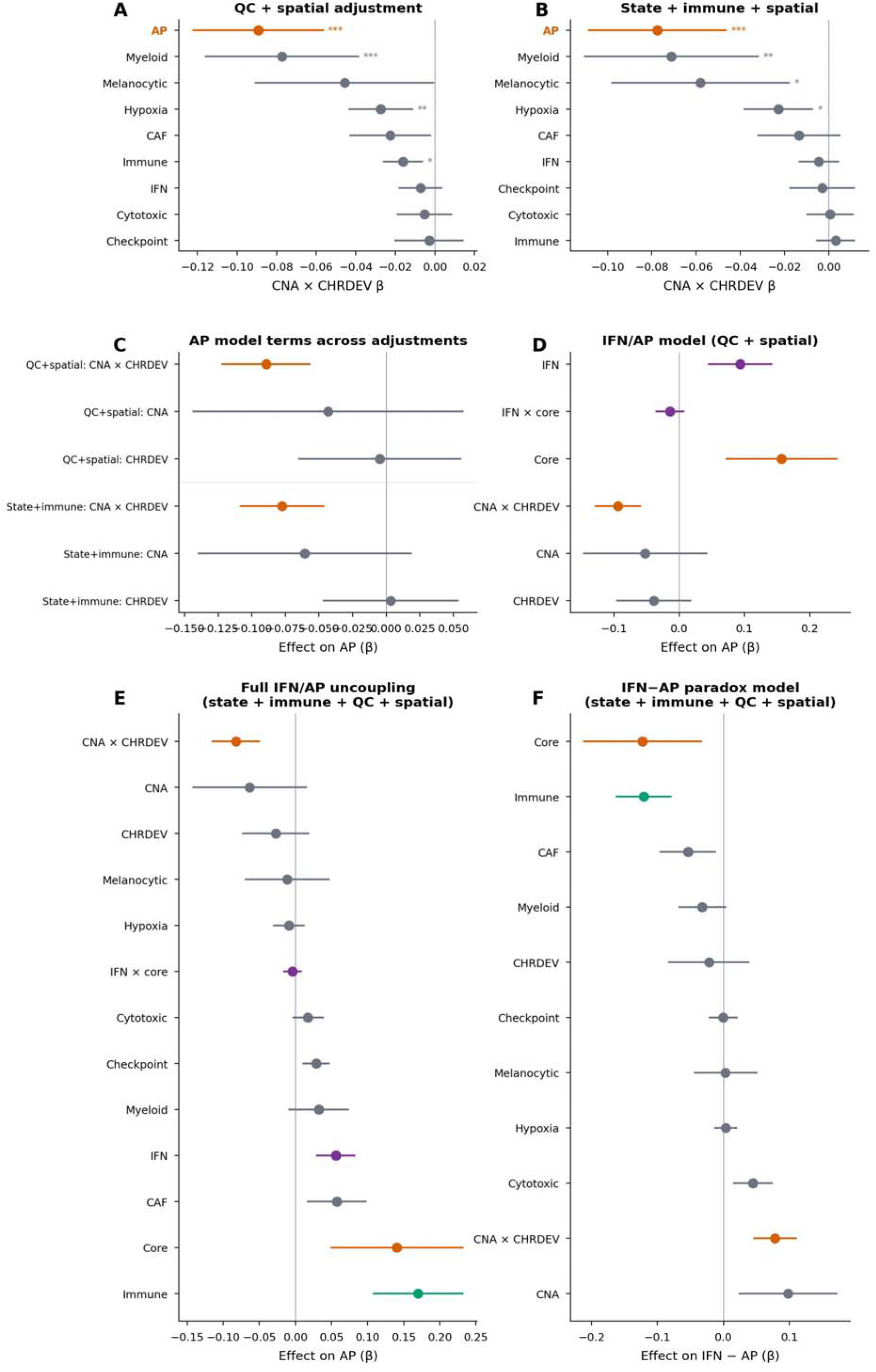
Full program specificity and IFN/AP uncoupling models

**Supplementary Figure 5:**
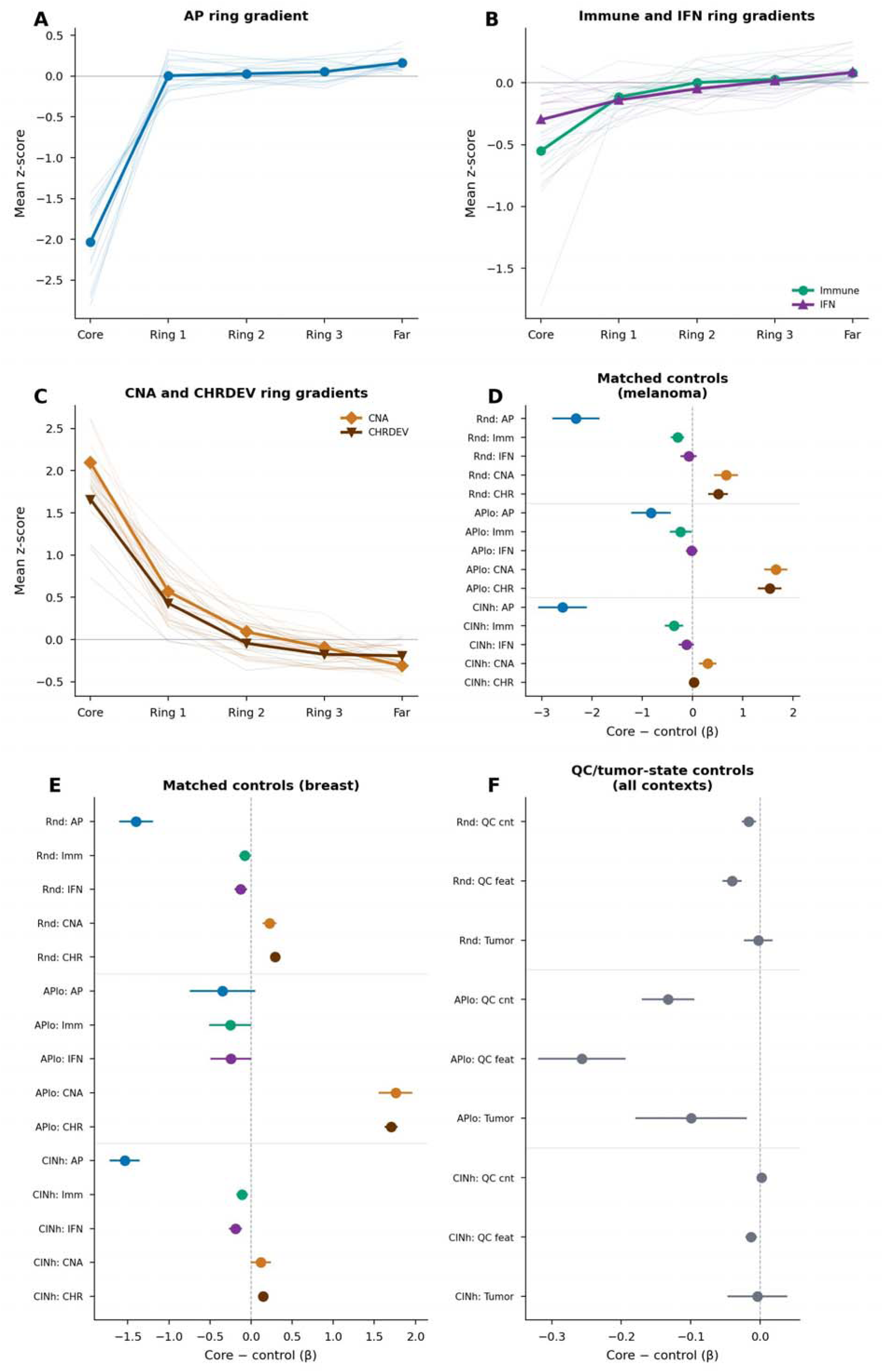
Ring gradients and matched controls across contexts

**Supplementary Figure 6:**
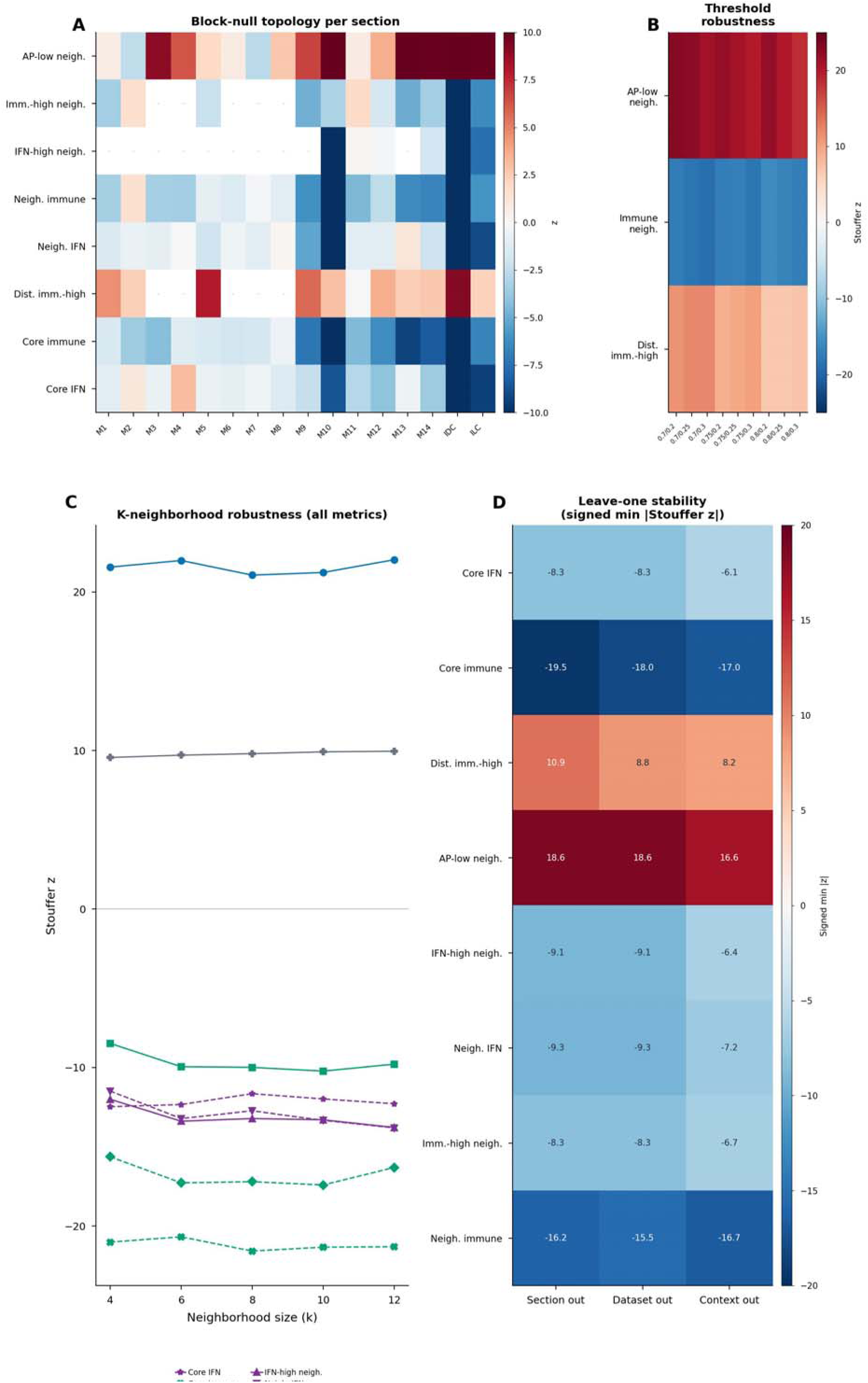
Full robustness - block-null, threshold, k-size, and leave-one stability

**Supplementary Figure 7:**
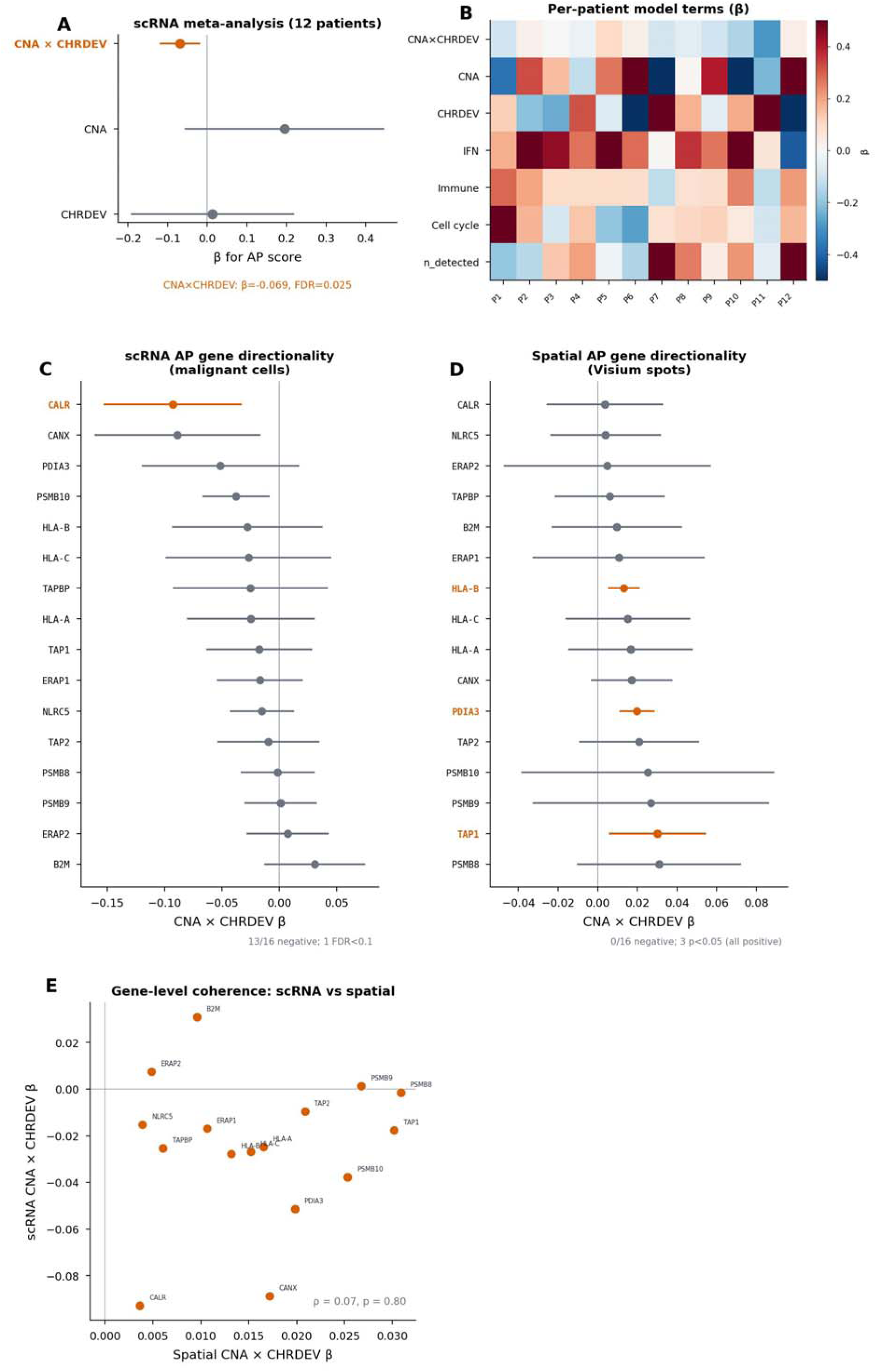
Malignant-cell scRNA validation and AP gene-level effects

**Supplementary Figure 8:**
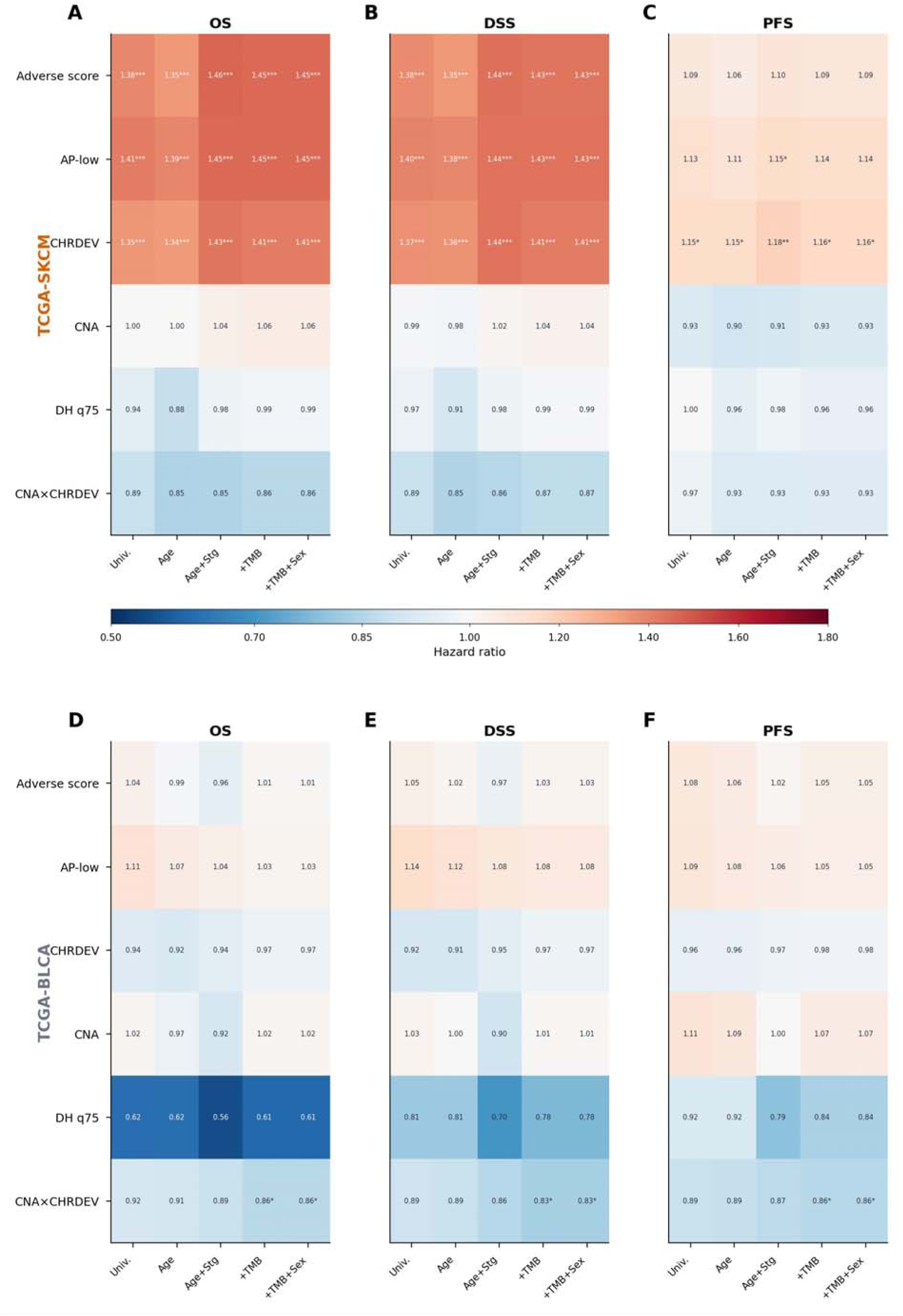
Full TCGA survival - all features, endpoints, and models

**Supplementary Figure 9:**
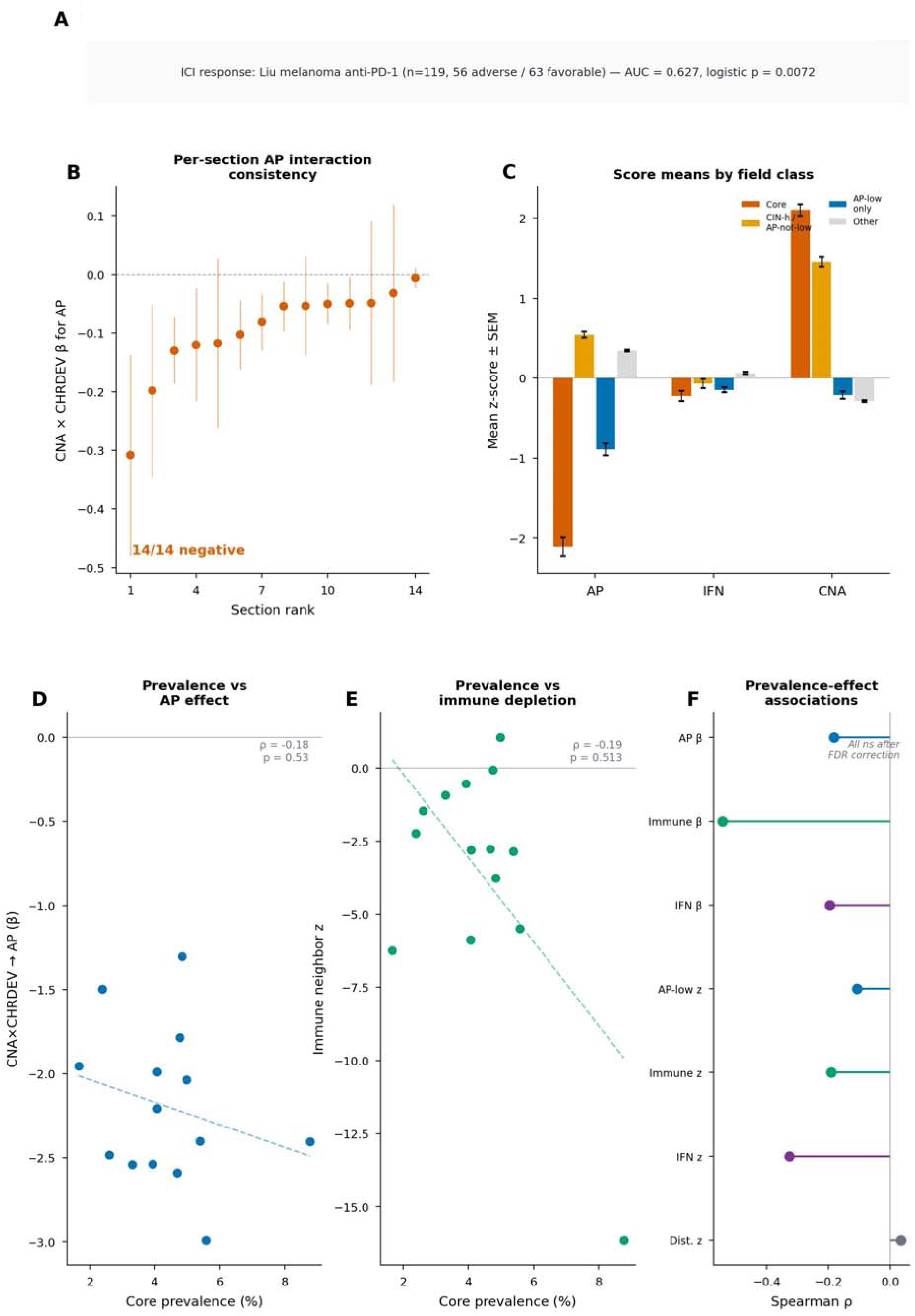
ICI projection, per-section consistency, and prevalence-effect

**Supplementary Figure 10:**
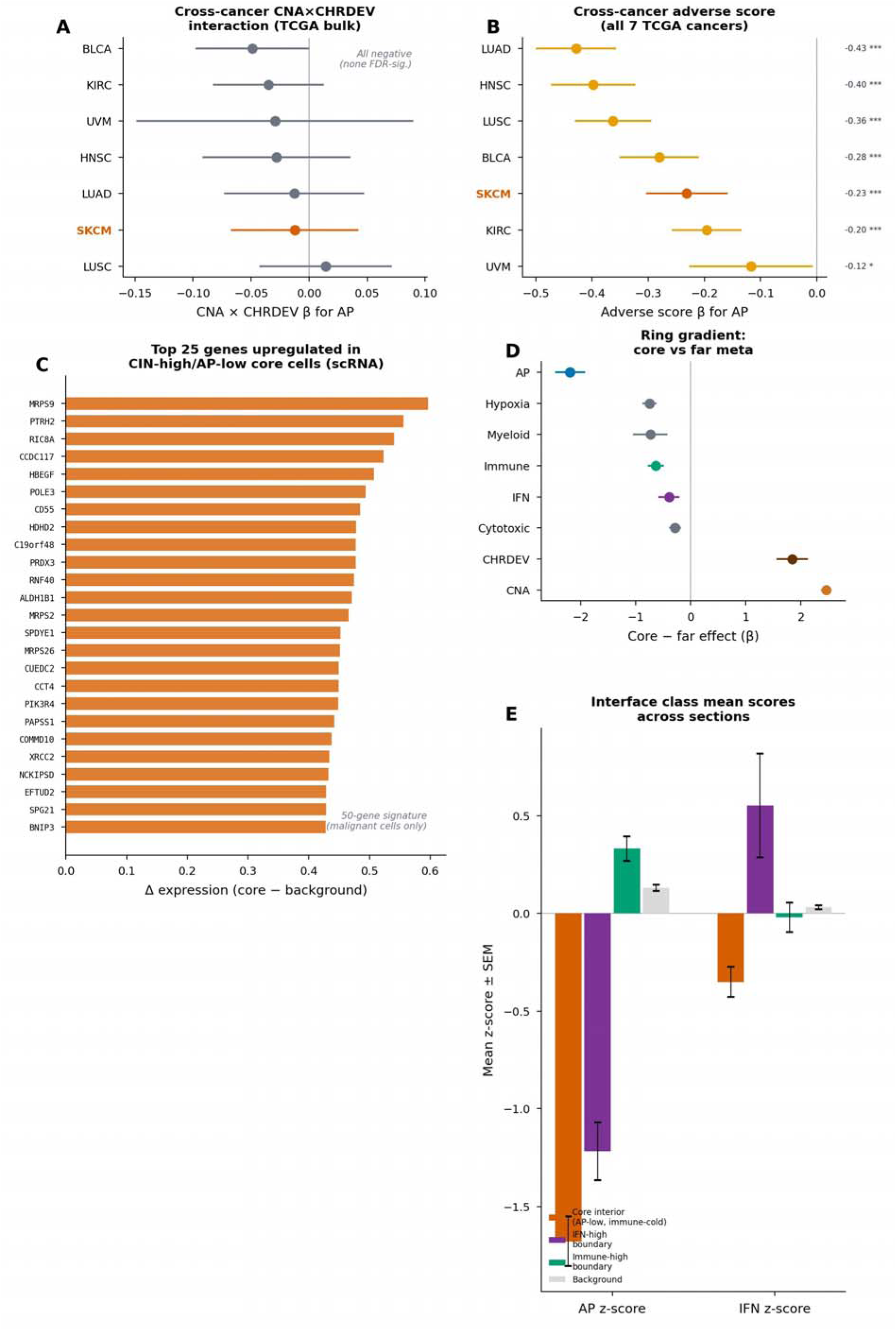
Cross-cancer generalizability, core gene signature, and spatial quantification

