## Supplementary File for "Transcriptome-inferred CIN-like fields mark antigen-presentation-low, immune-cold spatial neighborhoods"

Subhajit Dutta

Institute for Biochemistry and Molecular Cell Biology, Center for Experimental Medicine, University Medical Center Hamburg-Eppendorf, Hamburg, Germany

**This file contains:**

Supplementary Figure 1

Supplementary Figure 2

Supplementary Figure 3

Supplementary Figure 4

Supplementary Figure 5

Supplementary Figure 6

Supplementary Figure 7

Supplementary Figure 8

Supplementary Figure 9

Supplementary Figure 10

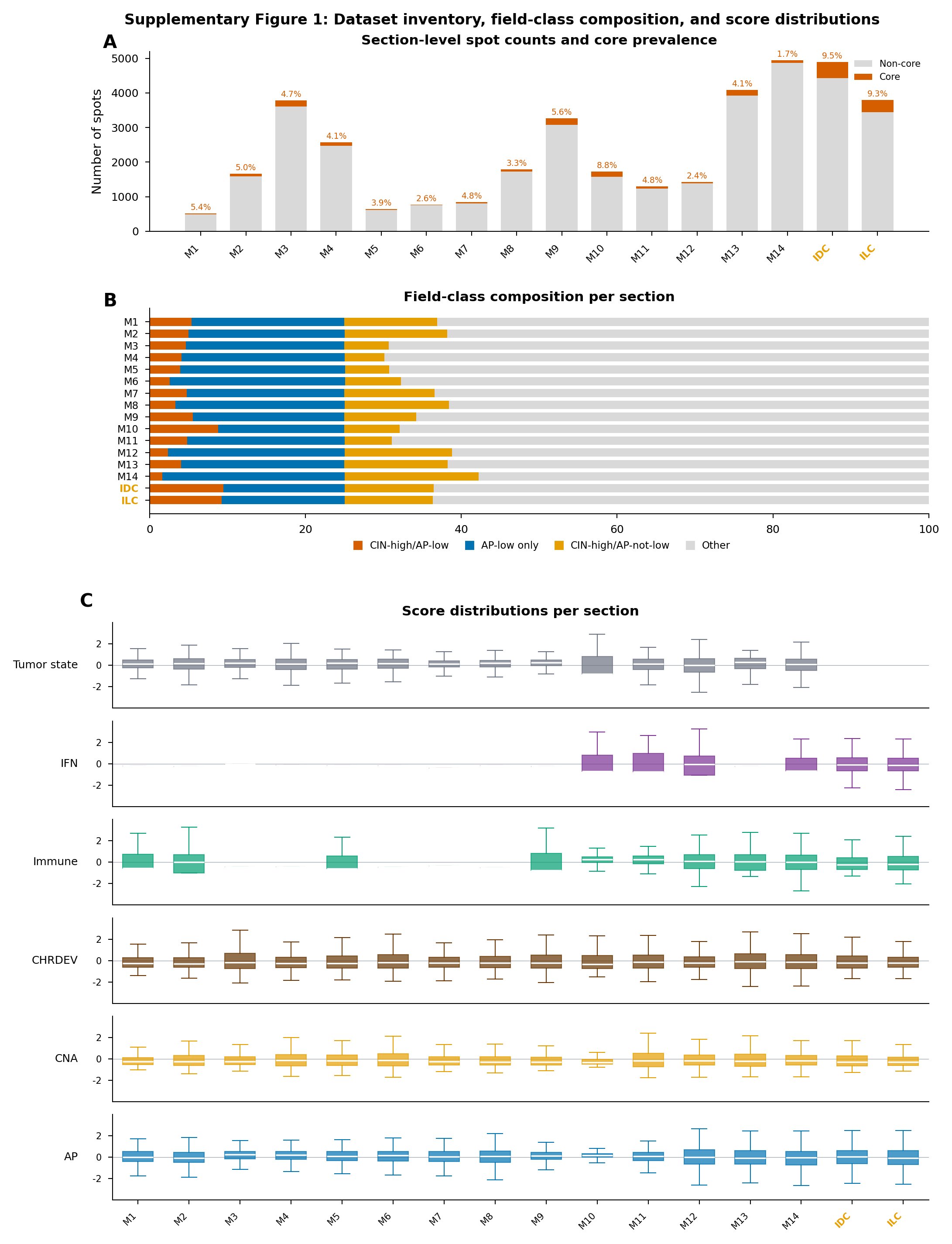

**Supplementary Figure 1 | Dataset inventory, field-class composition, and score distributions.** a, Section-level spot counts and CIN-high/AP-low core prevalence across 14 melanoma sections (M1–M14) and 2 breast Visium sections (IDC, ILC). Core prevalence (%) is annotated above each bar. b, Stacked bar chart of field-class composition per section, showing CIN-high/AP-low, AP-low only, CIN-high/AP-not-low and other fractions. c, Box-and-whisker plots of section-level score distributions for tumor state, IFN, immune, CHRDEV, CNA and AP z-scores. Flat IFN distributions in several GSE250636 sections (M1–M9) reflect limited IFN-pathway gene coverage in those datasets.

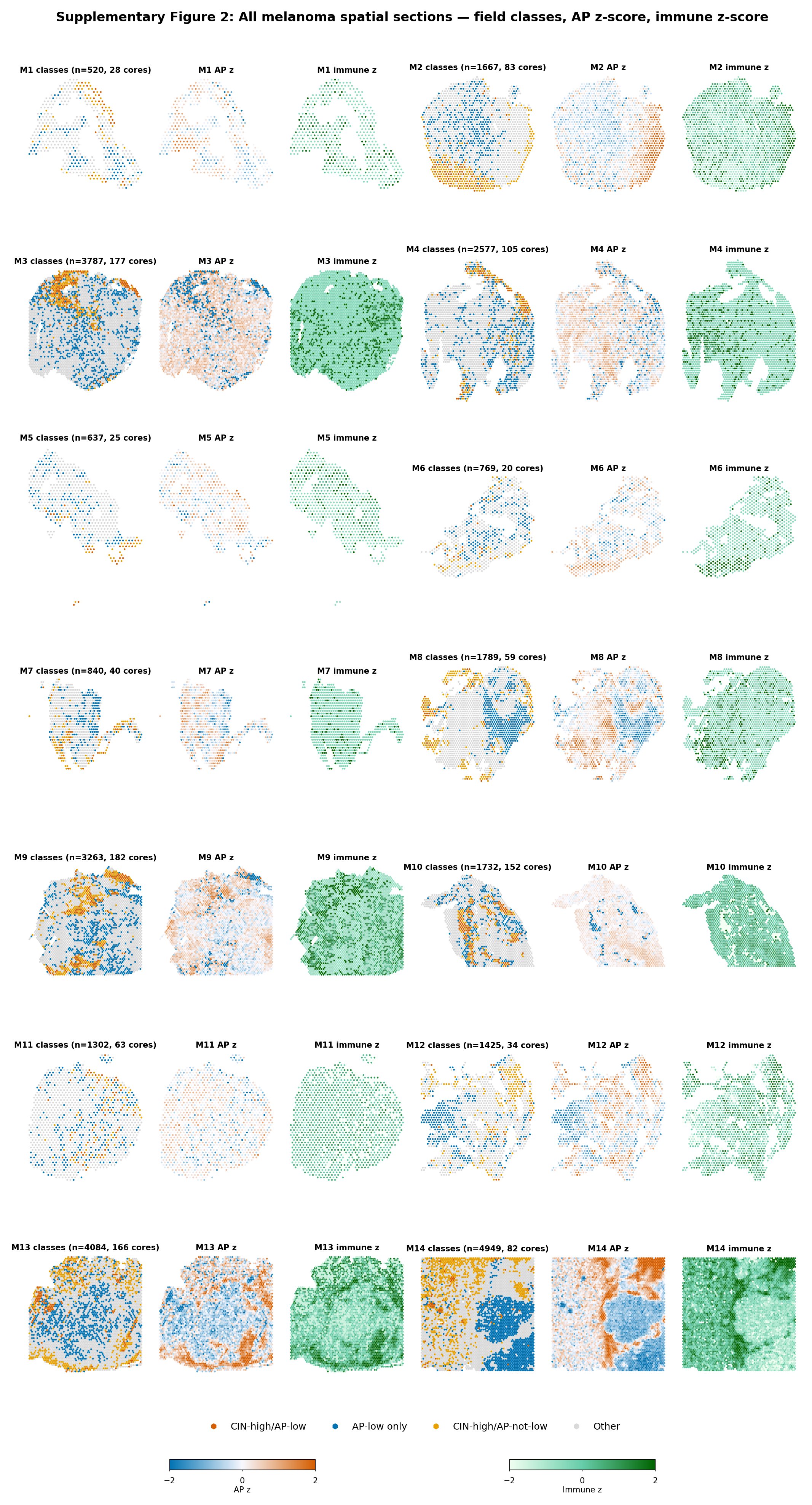

**Supplementary Figure 2 | Melanoma spatial atlas: representative field-class and score maps.** Spatial maps of field classes, CNA z-score, CHRDEV z-score, AP z-score, immune z-score and IFN z-score for all 14 melanoma tissue sections. Each row corresponds to one section (M1–M14). CIN-high/AP-low cores are shown in orange; AP-low-only spots in blue; CIN-high/AP-not-low spots in yellow; other spots in gray. Score maps use diverging color scales centered at zero.

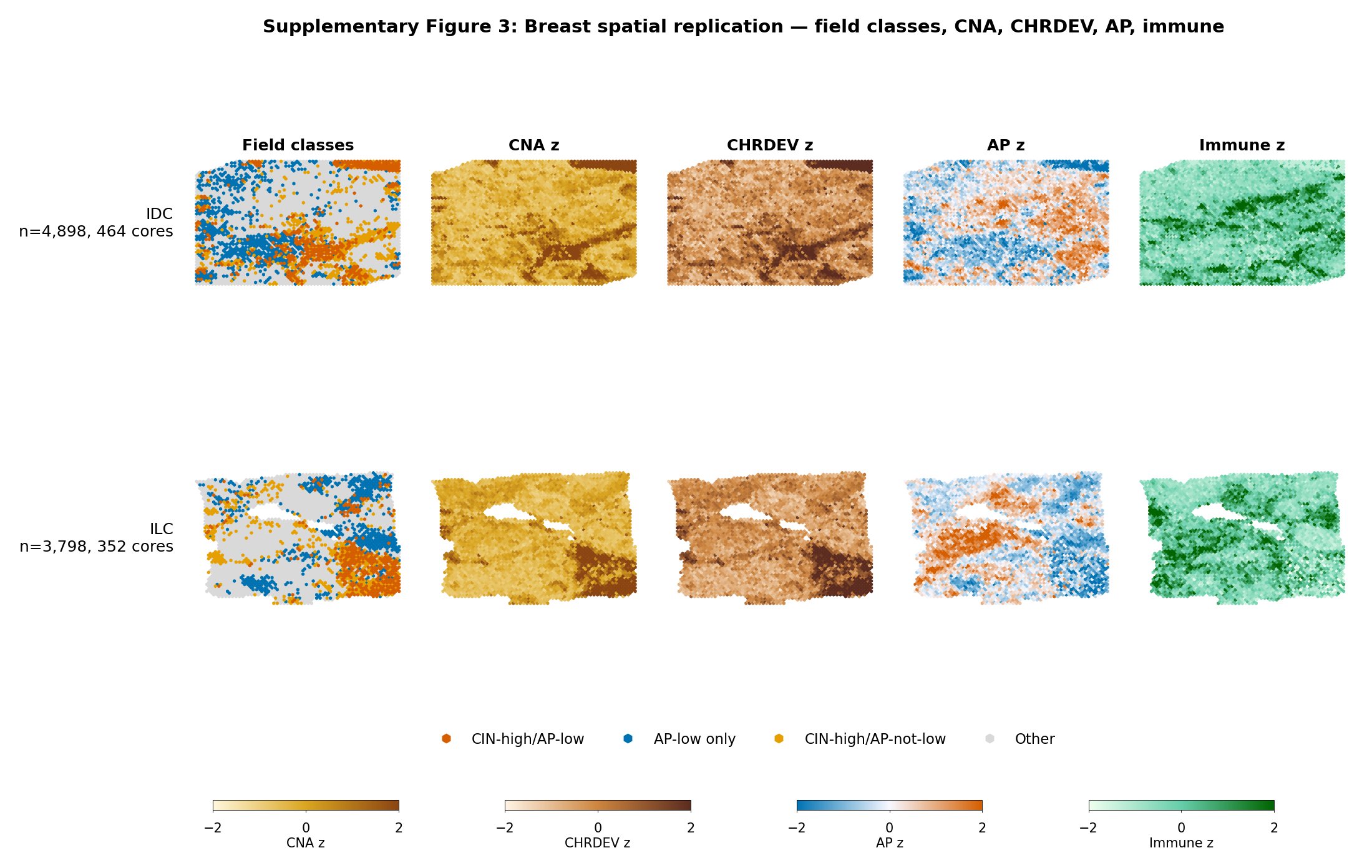

**Supplementary Figure 3 | Breast cancer Visium spatial atlas.** Spatial maps of field classes, CNA z-score, CHRDEV z-score, AP z-score, immune z-score and IFN z-score for the two breast Visium sections (IDC: ductal carcinoma in situ/invasive carcinoma, FFPE; ILC: Block A Section 1). Layout matches Supplementary Fig. 2.

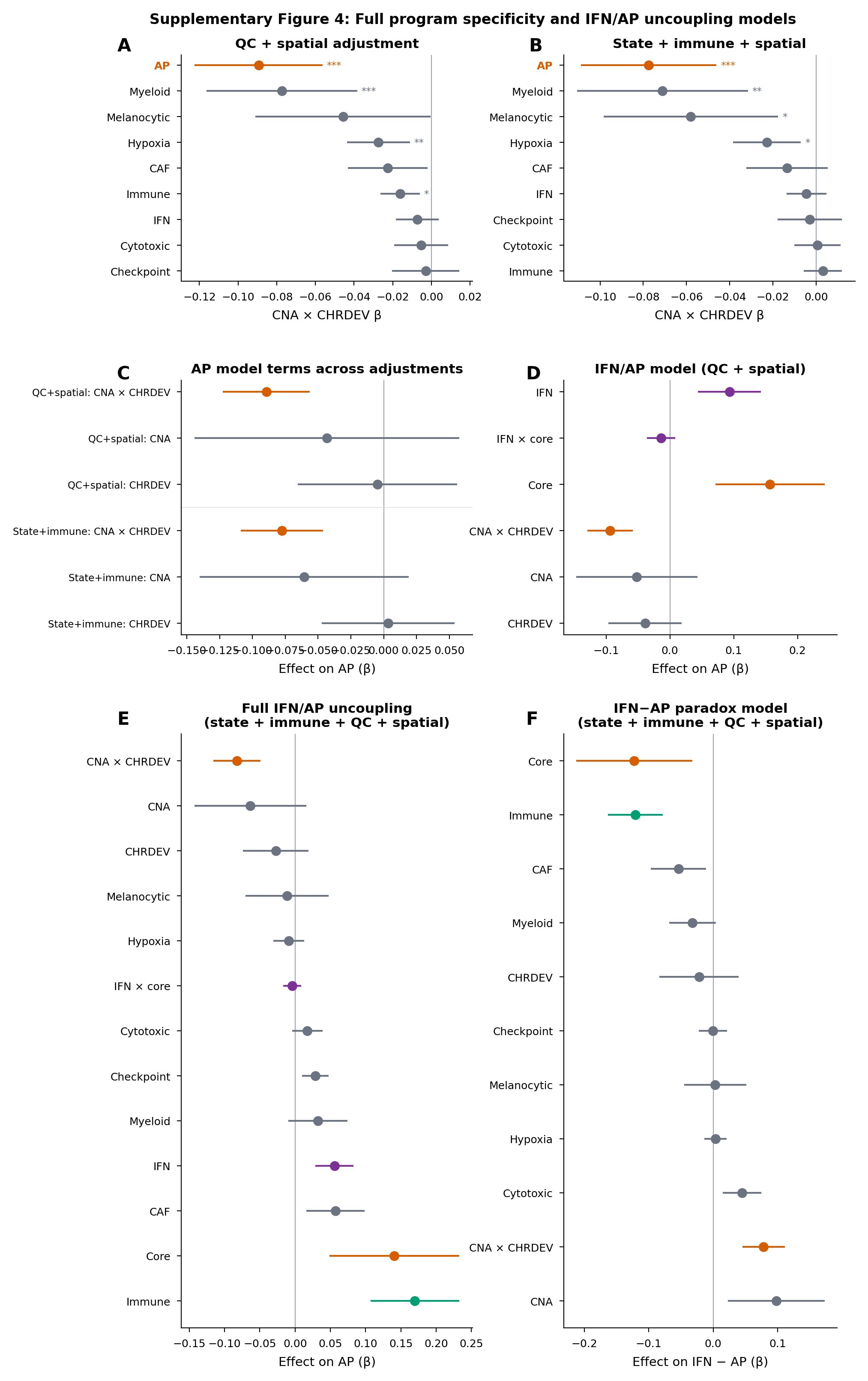

**Supplementary Figure 4 | Program-specific CNA-by-CHRDEV interaction effects.** a, Forest plots of the CNA-by-CHRDEV interaction coefficient for each program outcome (AP, immune, cytotoxic, checkpoint, IFN, CAF, hypoxia, melanocytic state, myeloid) from the section-level adjusted models. b, Section-level coefficients for each program, showing heterogeneity across melanoma sections. c, Covariate-adjustment sensitivity for non-AP programs. d, Leave-one-section stability for program-specific interaction effects.

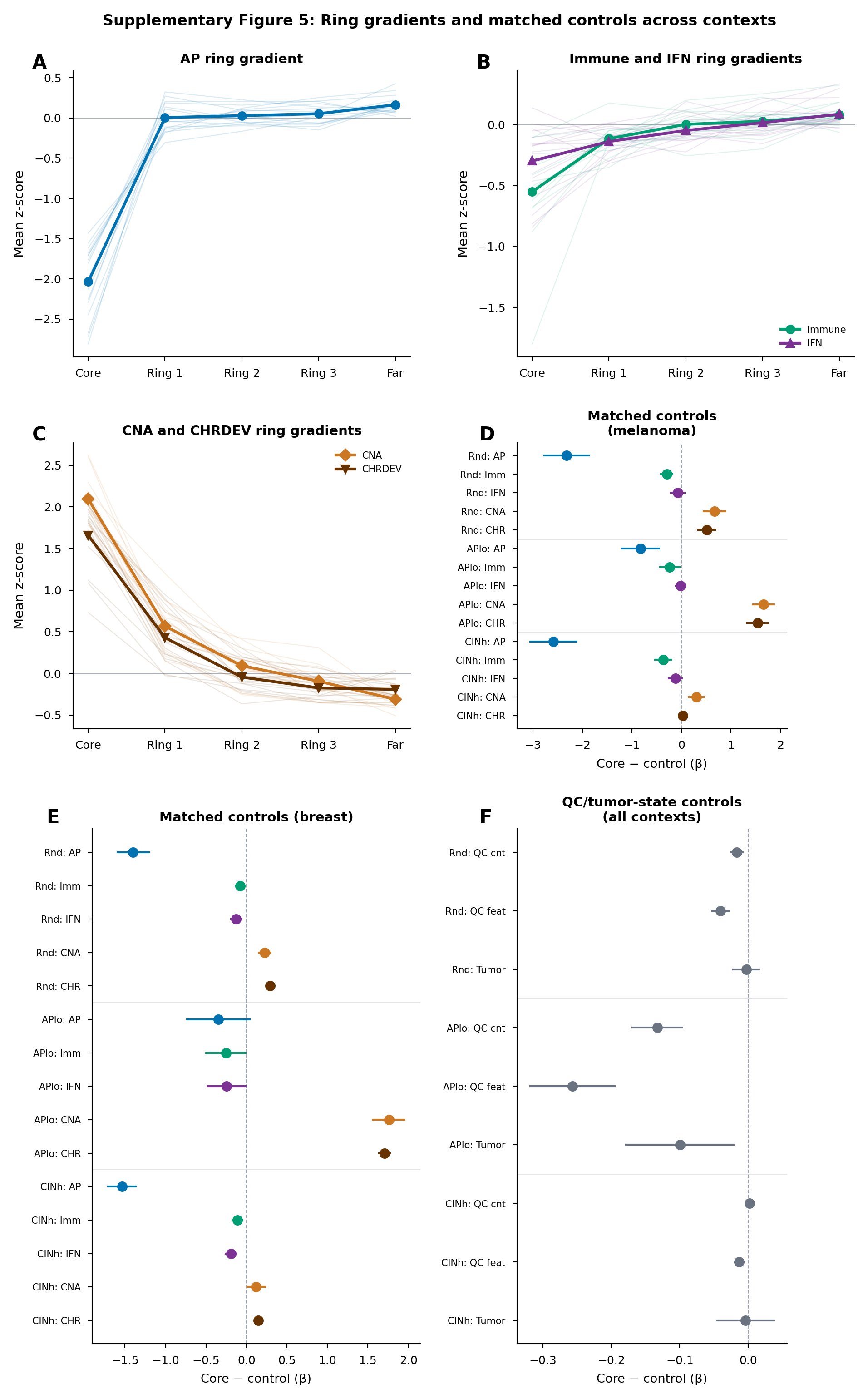

**Supplementary Figure 5 | Ring-gradient profiles and interface quantification.** a, Ring-gradient profiles around CIN-high/AP-low cores for AP, immune, IFN, CNA, CHRDEV, hypoxia, myeloid and cytotoxic programs across melanoma and breast sections. Each line represents one section; meta-analytic core-minus-far contrasts are annotated. b, Section-level interface-class mean scores for AP, immune and IFN across immune-cold AP-low interiors, IFN-high AP-low boundaries, immune-high boundaries and background regions. c, Ring-gradient meta-analytic summary with 95% confidence intervals.

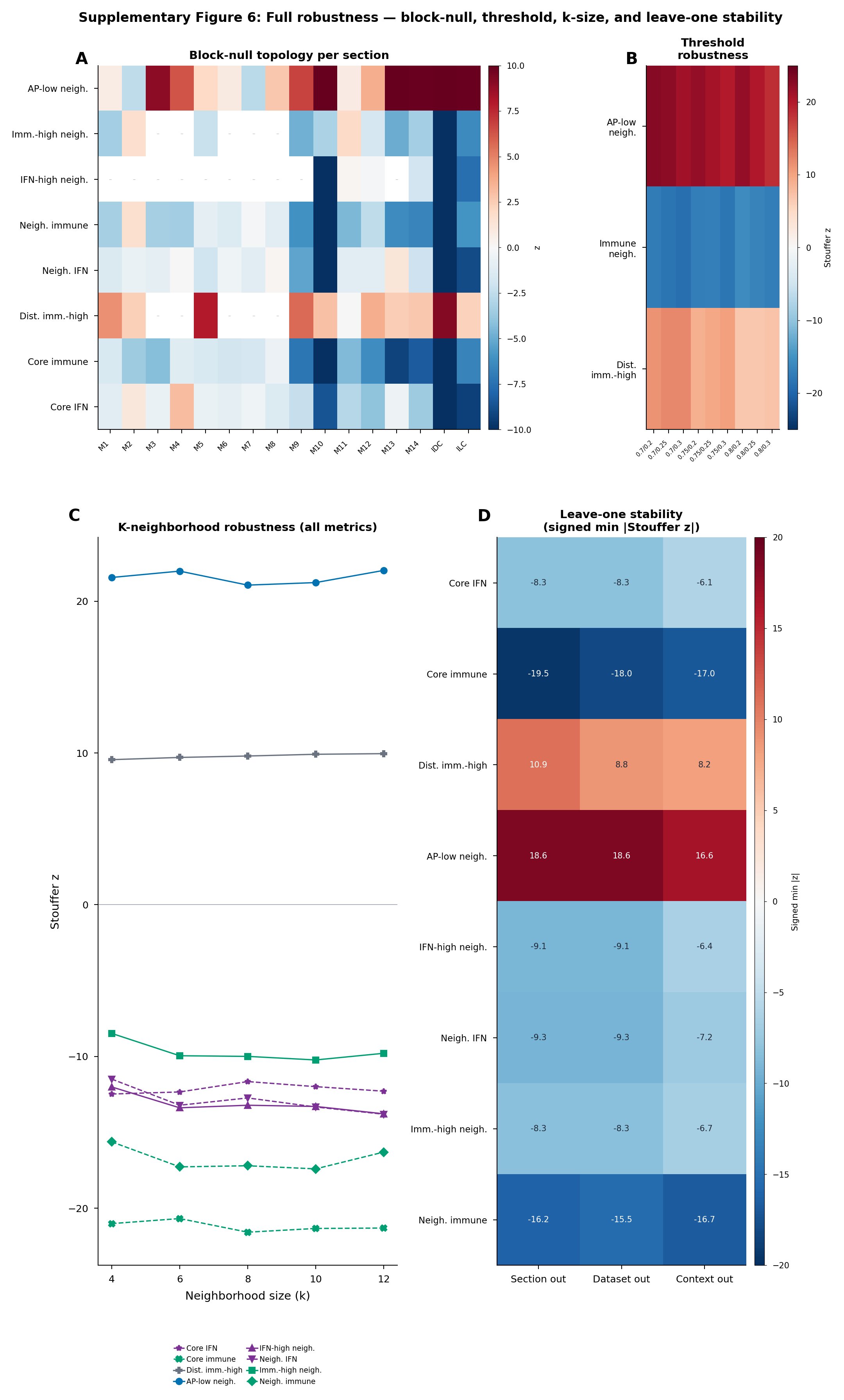

**Supplementary Figure 6 | Robustness and leave-one sensitivity analyses.** a, Leave-one-section stability for block-null topology metrics (AP-low neighbor enrichment, immune/IFN neighbor depletion, distance to immune-high). Each panel shows the Stouffer z when one section is removed. b, Leave-one-dataset stability, removing one GEO dataset at a time. c, Leave-one-context stability, removing melanoma or breast context. d, Threshold robustness across alternative CNA/CHRDEV quantile cutoffs and AP-low definitions. e, Neighborhood-size robustness across k = 4, 6, 8, 10 and 12 nearest neighbors.

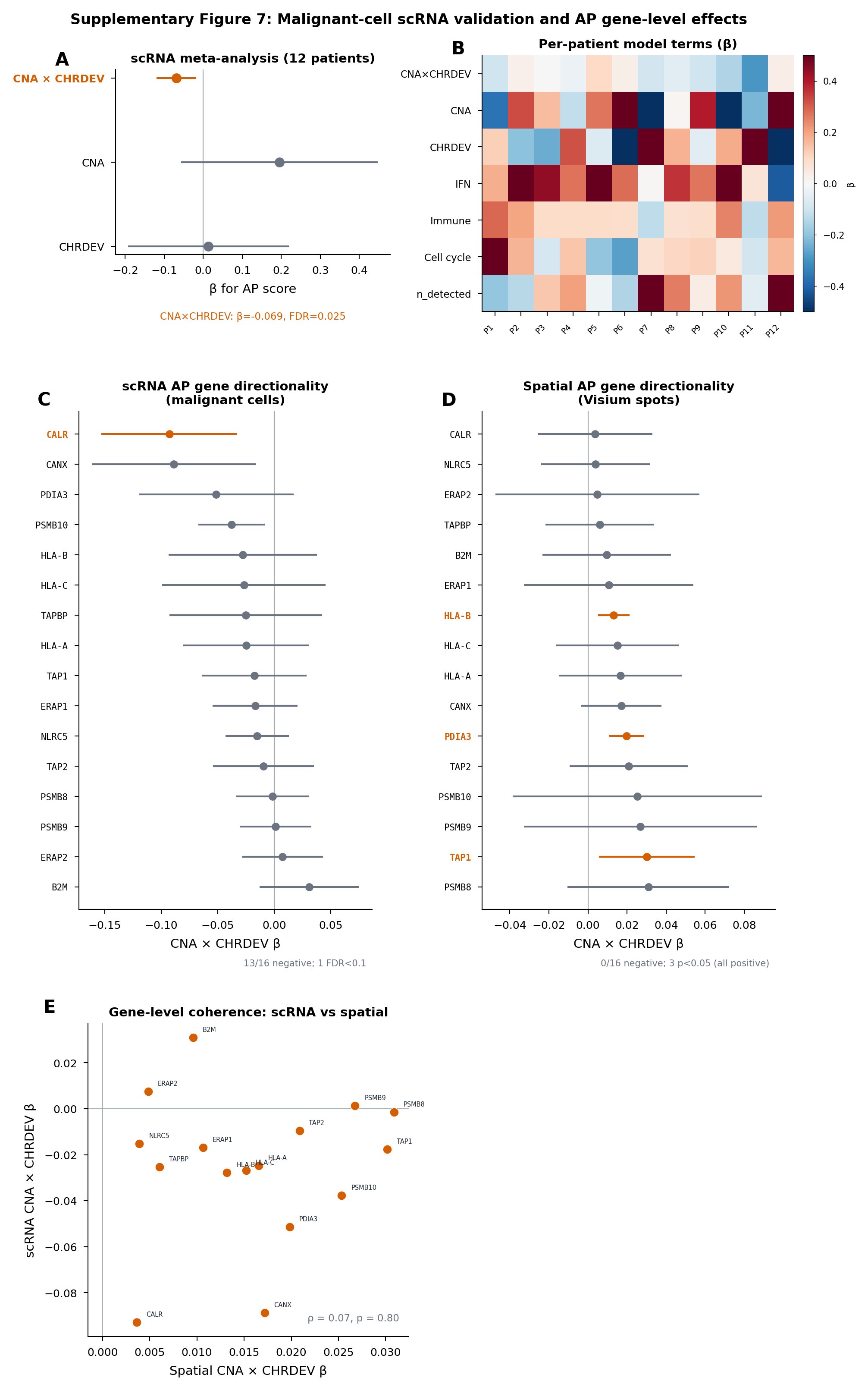

**Supplementary Figure 7 | Malignant-cell scRNA-seq validation and AP gene-level effects.** a, Random-effects meta-analysis of CNA-by-CHRDEV interaction, CNA main effect and CHRDEV main effect on AP activity across 12 melanoma patients (malignant cells only). b, Per-patient heatmap of model term coefficients (CNA×CHRDEV, CNA, CHRDEV, IFN, immune, cell cycle, n_detected) showing inter-patient heterogeneity. c, Gene-level CNA×CHRDEV coefficients for 16 AP machinery genes in malignant-cell scRNA-seq; 13/16 negative, 1 FDR < 0.1. d, Gene-level CNA×CHRDEV coefficients for the same 16 AP genes in spatial Visium spots; 0/16 negative, 3 significantly positive (p < 0.05), demonstrating that spot-level gene effects are confounded by spot composition and immune-cell AP expression. e, Scatter plot of spatial vs scRNA gene-level CNA×CHRDEV coefficients showing poor cross-platform coherence (ρ = 0.07, p = 0.80), supporting the pathway-level rather than gene-level interpretation.

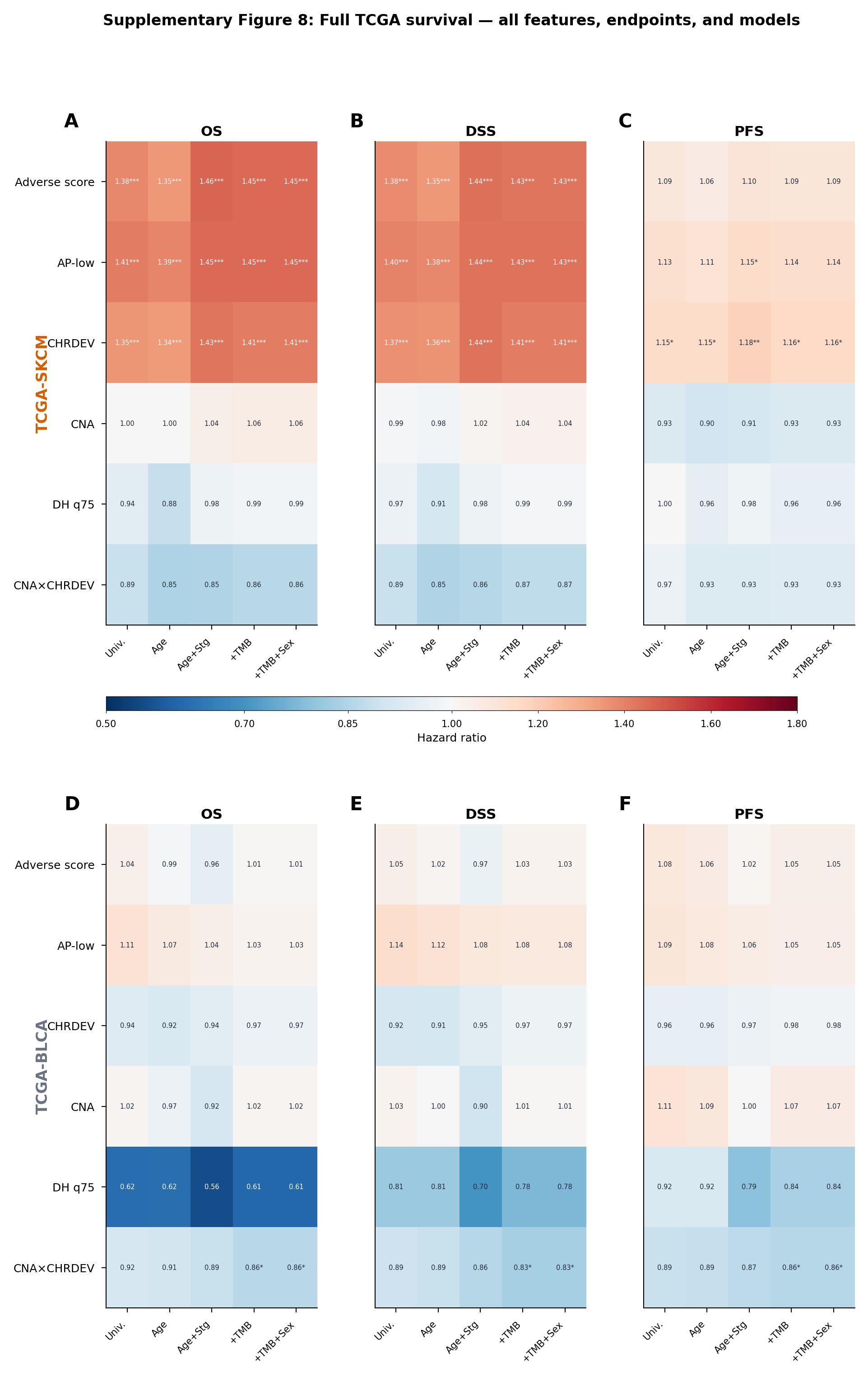

**Supplementary Figure 8 | TCGA survival analyses and feature-level models.** a, TCGA-SKCM Kaplan–Meier overall-survival curves stratified by adverse genotype-ecotype score (high vs low). b, Forest plot of hazard ratios for individual features (adverse score, AP-low score, CHRDEV score, CNA score, double-high q75, CNA×CHRDEV interaction) in TCGA-SKCM with age and stage adjustment. c, TCGA-BLCA overall-survival, disease-specific survival and progression-free survival hazard ratios for the adverse score with age and stage adjustment. d, TCGA-BLCA feature-level survival models, showing the absence of a consistent adverse-score signal in bladder cancer.

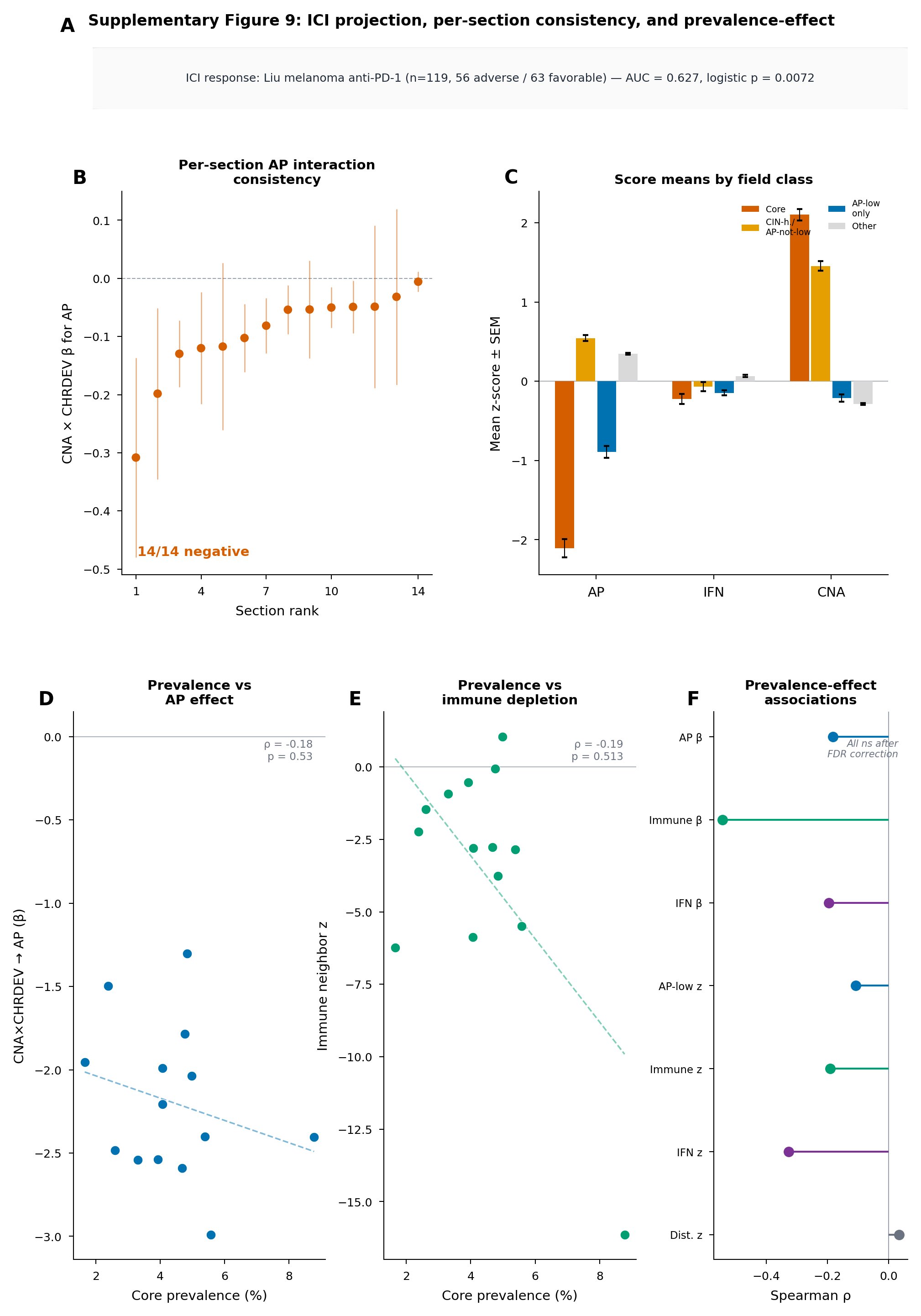

**Supplementary Figure 9 | Immunotherapy response projection.** a, Receiver operating characteristic (ROC) curve for the adverse genotype-ecotype score as a predictor of ICI non-response in melanoma cohorts (AUC = 0.627, logistic p = 0.0072, FDR = 0.072). b, Box plots of adverse-score distributions in ICI responders versus non-responders. c, Feature-level ICI-response associations for individual genotype-ecotype components. The result is presented as a projection test rather than a validated predictive classifier.

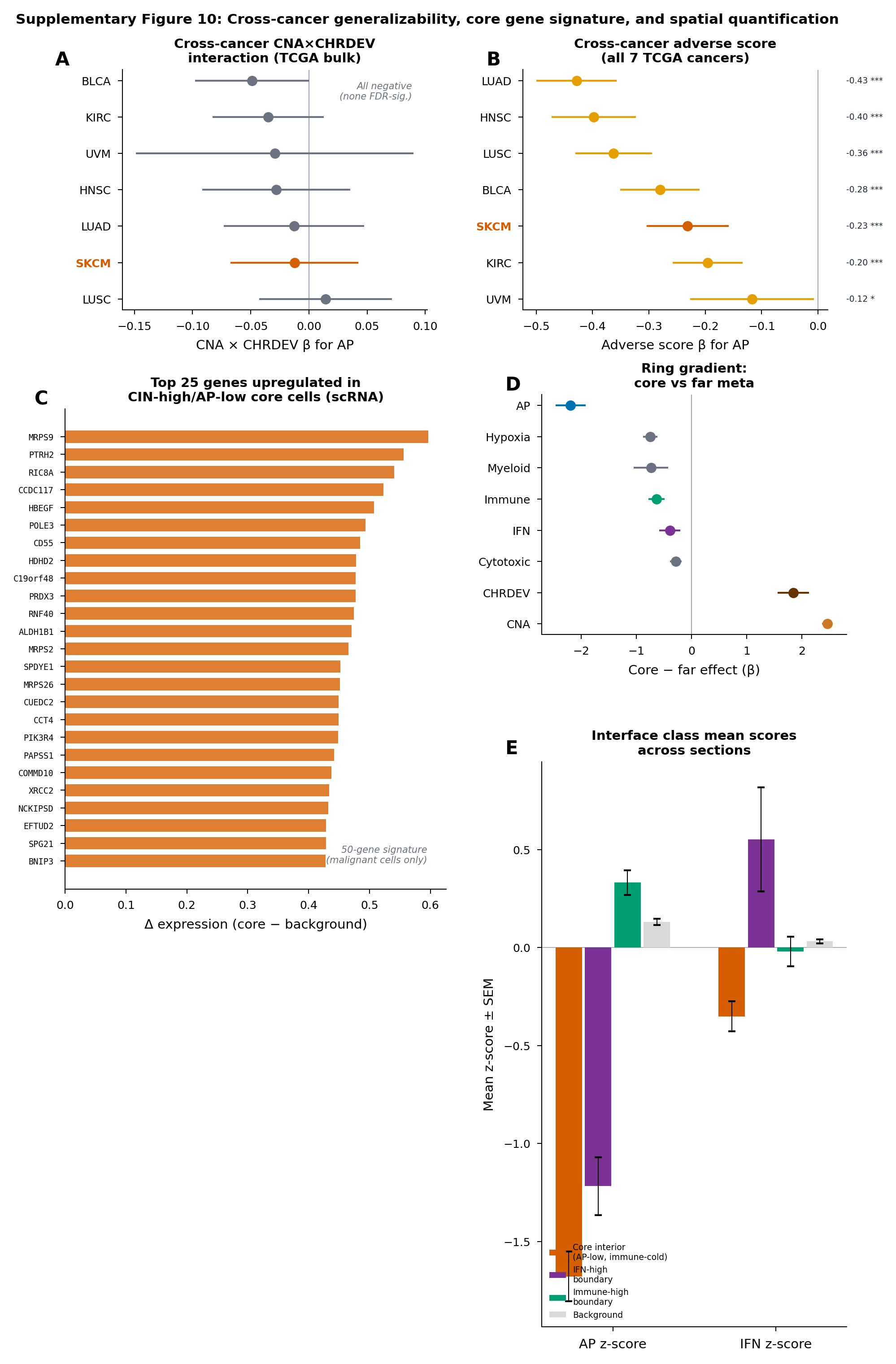

**Supplementary Figure 10 | Cross-cancer generalizability, core gene signature, and spatial quantification.** a, Cross-cancer CNA×CHRDEV interaction coefficients for AP in seven TCGA bulk RNA-seq cohorts (SKCM, LUAD, LUSC, HNSC, UVM, KIRC, BLCA). All directions negative; none FDR-significant. b, Cross-cancer adverse-score coefficients for AP across the same seven TCGA cancers, all significantly negative. c, Top 25 genes upregulated in CIN-high/AP-low core malignant cells relative to background malignant cells (from a 50-gene signature; scRNA-seq data). d, Meta-analytic ring-gradient core-minus-far contrasts for AP, hypoxia, myeloid, immune, IFN, cytotoxic, CHRDEV and CNA programs. e, Interface-class mean AP and IFN z-scores across sections for core interiors, IFN-high boundaries, immune-high boundaries and background regions.
